# A FIT2–BLTP1 module sustains lipid droplet phosphatidic acid and restrains overgrowth

**DOI:** 10.64898/2026.09.30.755522

**Authors:** Qingzhu Chu, Tiantian Zhou, Zhaomin Zhou, Shuhan Zhang, Lin Deng, Wei-Ke Ji

**Affiliations:** Department of Biochemistry and Molecular Biology, School of Basic Medicine, Tongji Medical College, and State Key Laboratory for Diagnosis and Treatment of Severe Zoonotic Infectious Diseases, Huazhong University of Science and Technology, Wuhan, Hubei 430030, China; Shenzhen Bay Laboratory, Shenzhen, 518132, China; Cell Architecture Research Center, Huazhong University of Science and Technology, Wuhan, Hubei 430030, China

## Abstract

Lipid droplets (LDs) are assembled at discrete endoplasmic reticulum (ER) subdomains whose local phospholipid composition, rather than their protein content alone, determines whether a LD nucleates and how large it grows. Phosphatidic acid (PA) is the pivotal lipid of this environment, yet how PA reaches a forming LD has remained unknown. Here we show that the bridge-like lipid transfer protein BLTP1 is required to maintain PA in LD-associated membranes. BLTP1 depletion impaired LD formation and produced fewer but enlarged LDs across mammalian cell types, and lowered LD accumulation during adipogenic differentiation. At the ER, BLTP1 engaged the LD emergence factor FIT2 through its N-terminal transmembrane segment, an interaction that strengthened upon induction of LD formation and was strongly reduced by the disease-associated FIT2 G232R substitution. Co-depletion of FIT2 relieved the LD overgrowth caused by loss of BLTP1. Wild-type BLTP1, but neither a FIT2-binding-defective chimera nor a lipid-transfer-groove mutant, rescued LD formation in BLTP1-depleted cells. Lipidomics of purified LDs showed that BLTP1 loss most strongly lowered PA, most prominently the species PA 36:2, whereas PA and the major phospholipid classes of bulk ER membranes were unchanged. Collectively, we propose that local PA supply by BLTP1 coordinates LD assembly and growth.

## Introduction

Lipid droplets (LDs) are the cellular stores of neutral lipids, with a triacylglycerol (TAG) and sterol-ester core enclosed by a phospholipid monolayer, and are the only organelles built de novo from the endoplasmic reticulum (ER) bilayer rather than by fission of a pre-existing compartment^1–3^. TAG synthesized within the bilayer first demixes into a nascent lens, which grows past a critical size, buds toward the cytosol and expands as further neutral lipid and monolayer phospholipids are delivered^1,3^. Each step is governed by membrane physics as much as by protein activity. The barrier to lens nucleation, the shape the lens adopts and the side of the bilayer from which the LD emerges are all set by the surrounding phospholipid composition, acting through surface tension, the intrinsic curvature of lipids such as phosphatidic acid (PA) and diacylglycerol, and asymmetry between the leaflets^4–7^. Consistent with this, LDs arise repeatedly at discrete ER subdomains that can be visualized before a LD becomes optically resolvable^8–10^. A LD assembly site is therefore defined not only by the proteins that mark it but also by the lipids that surround them.

The best-defined protein determinant of these sites is Seipin, the BSCL2 gene product mutated in Berardinelli–Seip congenital lipodystrophy^11^. Seipin concentrates at ER–LD junctions^12^, converts nascent lenses into mature LDs^10^, and forms an oligomeric cage whose luminal domain and hydrophobic helices accommodate a growing droplet^13–15^. It binds anionic phospholipids including PA^16^, clusters TAG in the bilayer to lower the nucleation barrier^17^, acts with LDAF1/promethin^18,19^, and stabilizes the ER–LD contacts through which TAG flows while counteracting ripening^20,21^; adipogenin remodels the complex to drive adipocyte growth^22^. This apparatus nucleates LDs but does not by itself specify the lipid composition of the membrane in which nucleation occurs.

The lipids themselves provide a second layer of control, and their origin matters. Localized recruitment of CTP:phosphocholine cytidylyltransferase stimulates phosphatidylcholine synthesis during LD expansion, whereas limiting phosphatidylcholine favors coalescence^23^; conversely, accumulation of PA, the GPAT/AGPAT product that Lipins convert to diacylglycerol ahead of DGAT-mediated TAG synthesis^24,25^, promotes fusion into supersized LDs^26^. Much of this chemistry is set at membrane contact sites by non-vesicular lipid transfer proteins^27^. ORP5 and ORP8 counter-transport phosphatidylserine and PI4P at ER contacts^28^. ORP5 occupies ER–LD contacts^29^, and ORP5/ORP8 recruit Seipin to PA-enriched mitochondria-associated ER subdomains^30^; Sec14L6 delivers phosphoinositide to LD-forming sites^31^. MIGA2 tethers mitochondria, ER and LDs^32^, and the bridge-like transporter VPS13A is found at ER–mitochondria–LD contacts^33^. Which lipid transfer protein, if any, supplies PA to a nascent LD site in mammalian cells has remained unknown.

FIT2 (*FITM2*) is the strongest candidate organizer of such an environment. FIT proteins are ER-resident, evolutionarily conserved proteins whose expression drives fat storage^34^; FIT2 has six transmembrane helices with cytosolic termini^35^, binds TAG directly^36^, and is required for nascent LDs to bud out of the ER rather than remain embedded or emerge lumenally^37^. FIT2 belongs to the lipid phosphatase/phosphotransferase superfamily^38^, acts as a fatty acyl-CoA diphosphatase that maintains ER structure and protects against ER stress^39,40^, and organizes LD biogenesis spatially by engaging ER tubule-forming proteins and septins^41^, placing it at the junction of phospholipid synthesis, surface tension and membrane curvature at emerging LDs^42,43^. Loss of FIT2 is consequential in vivo^44–47^, and biallelic *FITM2* variants cause a deafness–dystonia syndrome (Siddiqi syndrome)^48,49^. What FIT2 has lacked is a partner able to move phospholipid at the sites where it acts.

BLTP1 (KIAA1109/Csf1/Tweek/LPD-3) is such a protein. It is a founding member of the bridge-like lipid transfer protein family, built from tandem repeating β-groove (RBG) modules that form an elongated rod with a continuous hydrophobic channel^50–53^, the architecture established for VPS13 and ATG2, which transfer glycerophospholipids in bulk between tethered membranes^54–59^. Cryo-EM of the *C. elegans* orthologue LPD-3 revealed a rod-shaped, lipid-filled tunnel and identified its native partner subunits^60^; LPD-3 bridges the ER and plasma membrane^61^, and yeast Csf1 supplies phosphatidylethanolamine to the ER^62^. *Drosophila* Tweek maintains phosphoinositide homeostasis^63^, and TMEM170A is the associated ER scramblase^64^. Biallelic KIAA1109 variants cause Alkuraya–Kučinskas syndrome^65^, and Bltp1 loss is perinatally lethal in mice^66^, yet no mammalian membrane process has been assigned to BLTP1 at molecular resolution. In a companion study we identify the outer-mitochondrial-membrane protein FKBP8 as a partner of the BLTP1 C-terminal region^67^, the opposite end of the rod from the region examined here.

Here we identify BLTP1 as a determinant of LD biogenesis in mammalian cells. BLTP1 is required both for efficient LD formation and for restraining LD overgrowth, and it engages FIT2 through its N-terminal transmembrane (TM) segment, an interface weakened by the disease-associated FIT2 G232R substitution and strengthened upon induction of LD biogenesis. Suppressing FIT2 relieves the overgrowth caused by BLTP1 loss. Wild-type BLTP1, but neither a FIT2-binding-defective chimera nor a lipid-transfer-groove mutant, rescues LD formation in BLTP1-depleted cells. Lipidomics of purified LDs shows that BLTP1 loss selectively lowers PA, most prominently PA 36:2, whereas PA and the major phospholipid classes of bulk ER membranes are unchanged. We propose that local PA supply by BLTP1 determines how efficiently LDs nucleate and how large they grow.

## Results

### BLTP1 depletion produces fewer but much larger LDs

To ask whether the bridge-like lipid-transfer protein BLTP1^50,52,53^ contributes to LD biology, we depleted BLTP1 in U2OS cells with two independent siRNAs and imaged neutral lipid with BODIPY 558/568-C12 in live cells loaded with oleic acid (OA)^68^ (Fig. 1a). BLTP1-depleted cells contained fewer BODIPY-positive LDs than control cells (Fig. 1b), and the LDs that did form were much larger (Fig. 1c). Knockdown efficiency was confirmed by qPCR (Fig. 1d). The same fewer-but-larger pattern was seen in HeLa, Huh7 and HepG2 cells (Extended Data Fig. 1), indicating a general property of BLTP1 loss rather than a feature of one cell background. Both siRNAs produced the same result, and we used siRNA#3 in this study unless otherwise stated.

**Fig. 1.**
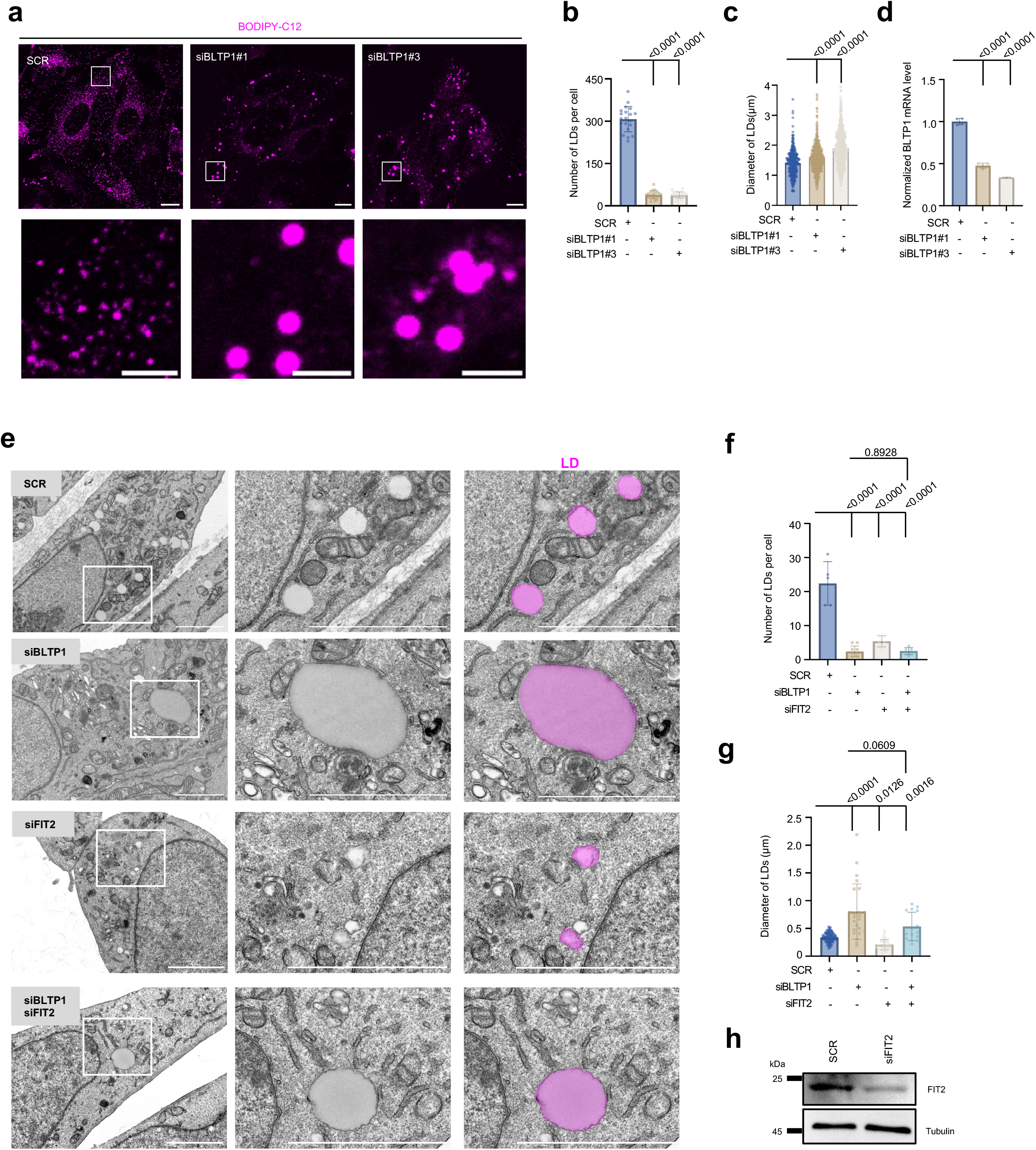
BLTP1 depletion results in fewer but larger lipid droplets. **a**, Representative live-cell confocal images of U2OS cells labeled with BODIPY 558/568-C12 after OA loading, transfected with non-targeting siRNA (SCR) or one of two independent BLTP1 siRNAs (siBLTP1#1, siBLTP1#3). **b**, LD number per cell and **c**, LD diameter quantified from **a** from at least three independent experiments. Mean ± SD. Ordinary one-way ANOVA with Tukey’s multiple-comparisons test. **d**, RT-qPCR quantification of BLTP1 mRNA, showing ∼50% (siBLTP1#1) and ∼65% (siBLTP1#3) reduction relative to SCR from three independent experiments. Mean ± SD. Ordinary one-way ANOVA with Tukey’s multiple-comparisons test. **e**, Representative transmission electron micrographs of U2OS cells treated with SCR, siBLTP1, siFIT2, or siBLTP1 + siFIT2 with ROIs to the right. **f**, LD number per cell section from **e**: SCR ≈22, siBLTP1 ≈2.4, siFIT2 ≈5.2, siBLTP1 + siFIT2 ≈2.3 (n = 5 cells per condition). Mean ± SD. Ordinary one-way ANOVA with Tukey’s multiple-comparisons test. **g**, LD diameter from **e**: SCR ≈0.30 μm, siBLTP1 ≈0.80 μm, siFIT2 ≈0.19 μm, siBLTP1 + siFIT2 ≈0.52 μm. Mean ± SD. Ordinary one-way ANOVA with Tukey’s multiple-comparisons test. **h**, FIT2 knockdown efficiency is validated by a representative immunoblot. Scale bar: 10 μm in whole-cell image and 2 μm in insets; 2 μm for TEM image in (e).

Transmission electron microscopy (TEM) consistently demonstrated that the number of LDs was significantly reduced in BLTP1-depleted cells compared to the control (Fig. 1e,f). Intriguingly, the residual LDs in BLTP1-depleted cells were greatly enlarged (Fig. 1e,g).

Interestingly, depletion of FIT2, an ER-resident acyl-CoA diphosphatase required for normal LD emergence from the bilayer^37,40^, also reduced LD number but, unlike BLTP1 depletion, reduced LD size (Fig. 1e–h). Simultaneous depletion of both proteins matched siBLTP1 in LD number, and its LD diameter was intermediate (≈0.52 µm, versus ≈0.80 µm with siBLTP1 alone and ≈0.30 µm with SCR; *P* = 0.061 versus siBLTP1; Fig. 1e–g), a trend consistent with FIT2 depletion alleviating the LD overgrowth in BLTP1-depleted cells.

Because FIT2 loss induces ER stress^40,46^, and because BLTP1 is an ER-anchored megaprotein whose depletion could perturb ER membrane homeostasis^61,62^, we asked whether the LD phenotype was an indirect consequence of a stressed or remodeled ER. Neither BLTP1 siRNA consistently raised the basal level of BiP, an ER stress marker, in the absence or presence of thapsigargin treatments (Extended Data Fig. 2a), and the frequency of ER whorls, a hallmark of ER stress, was not increased (Extended Data Fig. 2b,c). BLTP1 depletion therefore does not detectably activate the unfolded protein response under our conditions.

We next asked whether the reduced LD number reflected accelerated LD consumption rather than impaired formation. Contacts between LDs and LC3B-positive structures, and between LDs and LysoTracker-positive compartments, were largely unchanged by BLTP1 depletion (Extended Data Fig. 3a-c). Using a tandem RFP–GFP reporter fused to the LD coat protein PLIN2^69^, the number of acidified (RFP-only) puncta after 6 h of serum starvation was modestly lower, not higher, in BLTP1-depleted cells (Extended Data Fig. 3d,e). Increased lipophagic turnover therefore does not account for the loss of LDs.

### BLTP1 is required for efficient LD formation after OA loading

To investigate a potential role of BLTP1 in nascent LD formation, we delipidated cells and then triggered synchronous LD formation with OA, following the population over 60 min (Fig. 2a). Control cells accumulated LDs progressively, reaching ∼77 LDs per cell at 60 min, whereas BLTP1-depleted cells reached only ∼22 (Fig. 2b,c). Over the same interval, LDs in BLTP1-depleted cells grew larger than controls (Fig. 2d), so the fatty-acid load was not simply failing to be esterified but was instead partitioned into fewer, larger LDs. BLTP1 depletion therefore impairs LD biogenesis itself, a defect most consistent with impaired budding of nascent droplets from the ER bilayer, rather than a failure to synthesize neutral lipid.

**Fig. 2.**
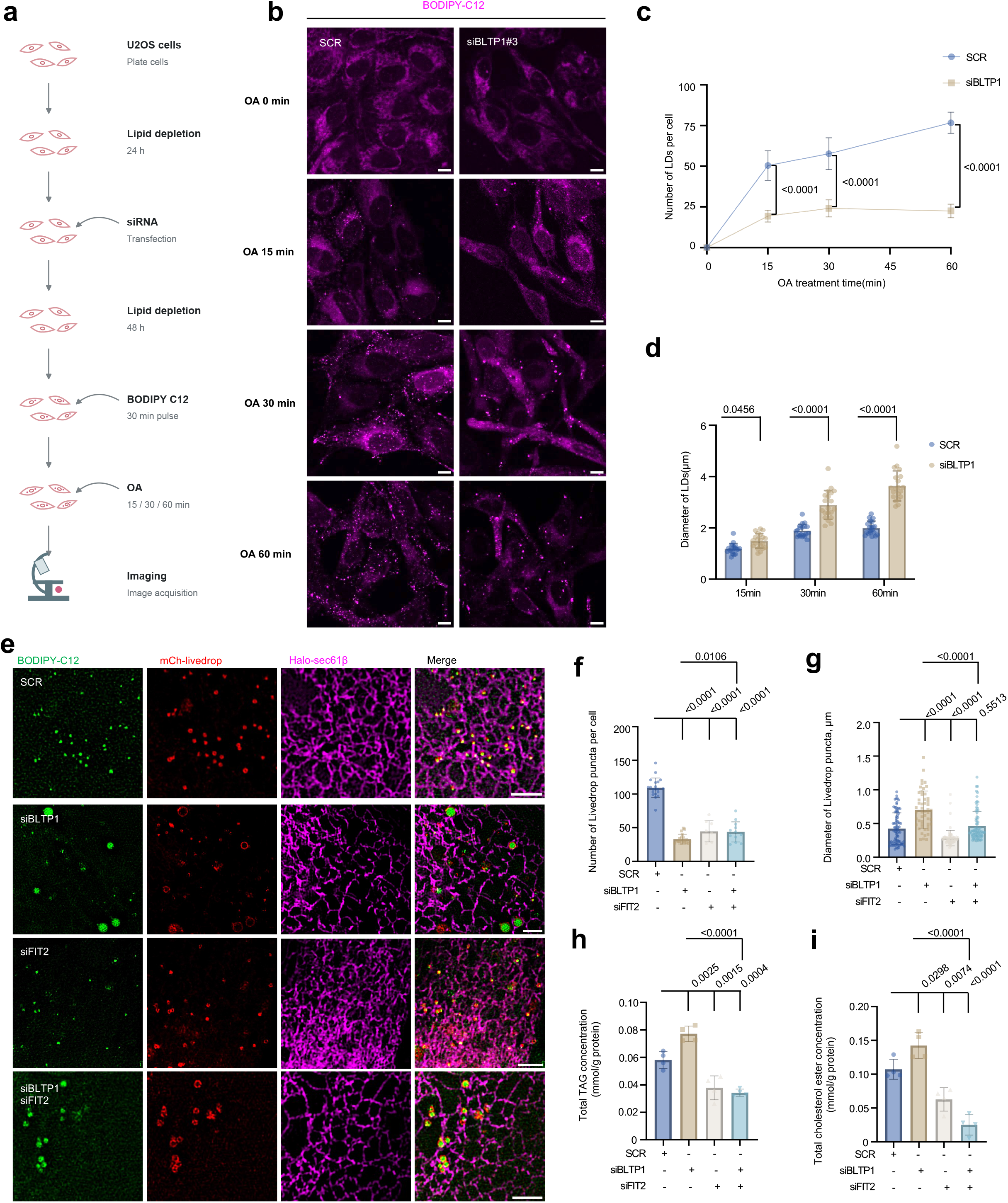
BLTP1 is required for nascent LD formation. **a**, Schematic of the de-lipidation and OA re-loading time course. **b**, Live BODIPY images of SCR and siBLTP1 U2OS cells at 0, 15, 30 and 60 min after OA addition. **c**, LD number per cell across the time course from three independent experiments. Mean ± SD. Ordinary one-way ANOVA with Tukey’s multiple-comparisons test. **d**, LD diameter at 15, 30 and 60 min from three independent experiments. Mean ± SD. Two-tailed unpaired Student’s t-tests. **e**, Live-cell 3D-SIM images of cells co-expressing mCh–LiveDrop (the GPAT4 amphipathic hairpin)^8,10^, and Halo–Sec61β labeled with JF646^70,71^, together with BODIPY, in SCR, siBLTP1, siFIT2 and siBLTP1 + siFIT2 cells. **f**, LiveDrop puncta per cell and **g**, LiveDrop puncta diameter from three independent experiments. Mean ± SD. Ordinary one-way ANOVA with Tukey’s multiple-comparisons test. **h**,**i**, Triacylglycerol (TAG; **h**) and total cholesterol ester (**i**) per unit protein in lysates of cells treated as in **Extended Data Fig. 4a** from four independent experiments. Mean ± SD. Ordinary one-way ANOVA with Tukey’s multiple-comparisons test. Scale bar: 10 μm in whole-cell image and 2 μm in insets.

We next visualized nascent LDs with mCherry-LiveDrop, the GPAT4 amphipathic hairpin that marks ER TAG lenses and newly forming droplets^8,10^, together with Halo-Sec61β to mark the ER^70,71^ and BODIPY-C12 to mark mature droplets, imaged live by super-resolution three-dimensional structured illumination microscopy (3D-SIM; Fig. 2e). In control cells, LiveDrop resolved into discrete puncta on the ER network. BLTP1 depletion reduced the number of LiveDrop puncta and increased their diameter, whereas FIT2 depletion reduced their number and decreased their diameter (Fig. 2f,g). In the double knockdown, puncta number stayed low, but puncta diameter returned to the control level (*P* = 0.55; Fig. 2f,g). Nascent LDs therefore behave like mature LDs: they are fewer and larger without BLTP1, fewer and smaller without FIT2, and co-depletion of FIT2 prevents the enlargement caused by BLTP1 loss. Importantly, in BLTP1-depleted cells, LiveDrop did not remain confined to small ER-embedded puncta but redistributed into rings surrounding the enlarged LDs, consistent with a nascent-site defect.

The two proteins act on the ER through entirely different mechanisms. FIT2 is a polytopic ER protein with acyl-CoA diphosphatase activity^38,40^, whereas BLTP1 is a repeating β-groove bridge predicted to carry glycerophospholipids^51,61^, and visualized in its nematode orthologue with ordered lipids filling a continuous channel^60^. We therefore compared single and combined knockdowns in the same experiment.

In U2OS cells loaded with OA, both single knockdowns reduced LD number to a similar degree, and co-depletion reduced it further; in contrast, in HepG2 cells the double knockdown gave no additional decrease in LD number (Extended Data Fig. 4a-f). LD size, however, moved in opposite directions. BLTP1 depletion approximately doubled mean diameter, whereas FIT2 depletion produced droplets slightly smaller than control (Extended Data Fig. 4a,c). In cells depleted of both proteins, mean diameter was intermediate, indicating that the overgrowth caused by BLTP1 loss was substantially suppressed by the additional loss of FIT2. The size phenotype was also reproduced in HepG2 cells (Extended Data Fig. 4d,f), and the suppression of the LD overgrowth induced by BLTP1 depletion by FIT2 loss is therefore not cell-line specific.

Because a reduction in LD number can arise either from fewer nucleation events or from a failure to make neutral lipid at all, we measured cellular lipid content directly. BLTP1 depletion did not lower triglyceride or cholesteryl ester; it raised both moderately (Fig. 2h,i). FIT2 depletion and the double knockdown each lowered both (Fig. 2h,i). Losing FIT2 therefore lowers neutral lipid and gives fewer, slightly smaller LDs, whereas losing BLTP1 raises triglyceride yet packages it into roughly four-fold fewer and two-fold wider LDs (Fig. 2c,d).

BLTP1 and FIT2 are both required at the nascent-droplet step, yet losing them moves LD size in opposite directions, and losing FIT2 relieves the overgrowth caused by losing BLTP1. That pattern has two explanations. The two proteins either work at different places or work at the same place at different times. We therefore asked whether they interact.

### BLTP1 directly interacts with FIT2

BLTP1 is a ∼5,000-residue RBG bridge whose rod is tethered to the ER by a single N-terminal transmembrane (TM) helix, with the remainder of the protein projecting into the cytosol as an elongated, lipid-filled tunnel^50–53,60,61^. Because the ER end of such a bridge may be positioned by a membrane-embedded partner, and because our loss-of-function data placed BLTP1 at the nascent-LD step alongside FIT2, we asked whether the two proteins interact with each other. GFP-Trap assays showed that BLTP1-GFP captured ectopic Halo-FIT2 (Extended Data Fig. 5a) and endogenous FIT2 (Extended Data Fig. 5b), but not the abundant ER membrane proteins RTN4 and ATL3 (Extended Data Fig. 5c).

To test the interaction on endogenous protein, we used a U2OS line in which the endogenous BLTP1 locus carries a C-terminal 3×Flag tag^67^. Flag-Trap immunoprecipitation from these cells pelleted endogenous FIT2, which was absent from precipitates of control cells expressing the empty 3×Flag-N1 vector, establishing that the two endogenous proteins indeed interact at native expression levels (Fig. 3a).

**Fig. 3.**
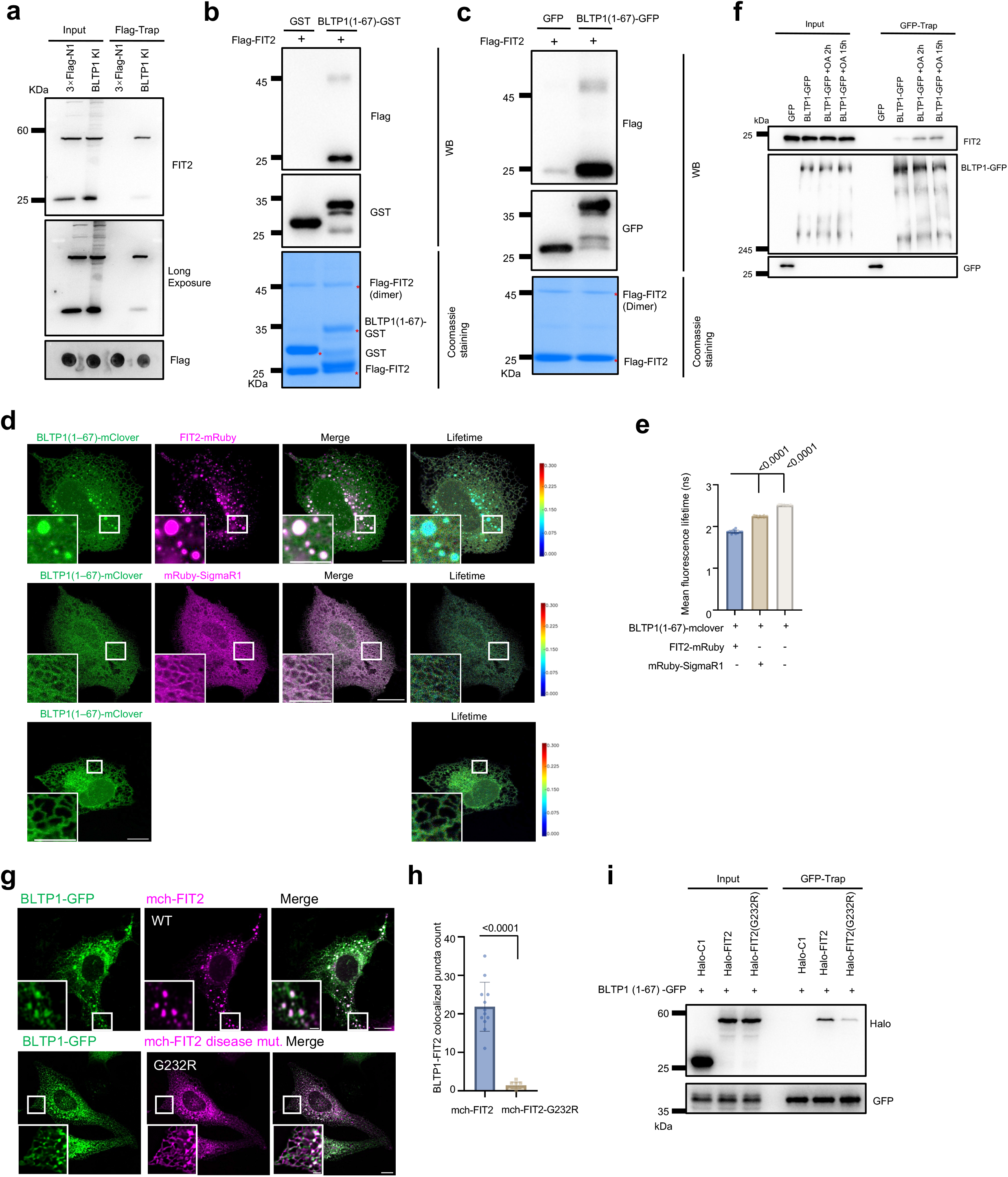
BLTP1 engages FIT2 through a transmembrane interface. **a**, Representative endogenous co-immunoprecipitation from the BLTP1-3×Flag knock-in U2OS line using Flag-Trap resin, immunoblotted for FIT2; cells expressing the empty 3×Flag-N1 vector serve as the negative control. Bottom, anti-Flag dot blot. **b**, Representative GST pull-down of purified Flag–FIT2 with BLTP1(1–67)–GST or GST alone, with the corresponding Coomassie-stained gel. **c**, Representative GFP-Trap pull-down of purified Flag–FIT2 with BLTP1(1–67)–GFP or GFP alone. **d**, Representative FLIM-FRET^72^ of BLTP1(1–67)–mClover as donor with FIT2–mRuby, mRuby–SigmaR1, or donor alone. **e**, mean donor fluorescence lifetime from three independent experiments. Mean ± SD. Ordinary one-way ANOVA with Tukey’s multiple-comparisons test. **f,** Representative GFP-Trap co-immunoprecipitation of endogenous FIT2 with GFP or BLTP1–GFP from cells left untreated or loaded with OA for 2 h or 15 h. **g**, Representative confocal imaging of U2OS cells expressing BLTP1–GFP (green) with wild type or the Siddiqi-syndrome variant FIT2(G232R)^49^ (magenta). **h**, Colocalized puncta per cell (262 puncta from 12 cells in case of mCh–FIT2 WT; 14 puncta from 10 cells in case of mCh-FIT2 G232R) from three independent experiments. Mean ± SD. Two-tailed unpaired Student’s t-tests. **i**, Representative GFP-Trap pull-down comparing binding of BLTP1(1–67)–GFP to Halo alone, Halo–FIT2 wild type and Halo–FIT2 G232R. Scale bars: 10 μm in whole-cell image and 2 μm in insets.

To test whether the interaction between BLTP1 and FIT2 is specific, we compared FIT2 with a panel of ER and LD-biogenesis proteins. Live-cell imaging showed that FIT2 overexpression caused ER membrane remodeling^41^, and BLTP1-GFP greatly overlapped with FIT2 at these ER structures (Extended Data Fig. 5d,e), but showed no comparable overlap with the tubule-shaping proteins RTN4a or ATL3, with ACSL3^9^, with LDAF1^18,19^, or with Seipin (BSCL2)^10,12^ (Extended Data Fig. 5f). Time-lapse imaging showed that these ER structures positive for BLTP1 and FIT2 were dynamic and underwent fusion (Extended Data Fig. 5e, bottom panel). BLTP1 therefore engages FIT2 selectively among the ER and LD-biogenesis proteins tested.

To map the FIT2-binding region, we expressed GFP-tagged BLTP1 fragments spanning the full protein (residues 1–67, 68–1500, 1501–2610, 2611–3592 and 3593–5093; Extended Data Fig. 6a). In cells, only the N-terminal fragment containing the transmembrane segment, BLTP1(1–67), colocalized with Halo–FIT2 in discrete puncta; none of the fragments covering the rest of the protein did (Extended Data Fig. 6b). Consistently, BLTP1(1–67)–GFP captured Halo–FIT2 in GFP-Trap assays (Extended Data Fig. 6c) and recovered substantially more Halo–FIT2 than did the transmembrane segment of calnexin (Calnexin-TM–GFP), an unrelated single-pass ER membrane protein (Extended Data Fig. 6d). The FIT2-binding determinant therefore resides within the short N-terminal, membrane-anchored segment of BLTP1, suggesting that FIT2 positions the ER-proximal end of the bridge rather than contacting the rod itself.

We next asked whether the association is direct. Recombinant BLTP1(1–67) fused to GST, but not GST alone, pelleted purified Flag-FIT2 in a pull-down assay, with input and bound proteins alongside a Coomassie-stained gel of the purified proteins (Fig. 3b). Consistently, GFP-Trap capture of BLTP1(1–67)–GFP, but not of GFP, pelleted purified Flag-FIT2 (Fig. 3c). These pull-downs indicate a direct interaction between the N-terminal TM region of BLTP1 and FIT2.

To test whether the interface forms in intact cells, we measured donor fluorescence lifetime by FLIM-FRET^72^. The mean lifetime of BLTP1(1–67)–mClover fell significantly when co-expressed with FIT2–mRuby, compared with the donor alone or with co-expression of the unrelated ER sheet membrane protein mRuby–SigmaR1^31^ (Fig. 3d,e). The lifetime reduction with FIT2 is larger than with a co-resident ER membrane protein at comparable expression, indicating closer or more frequent donor–acceptor apposition with FIT2 than can be accounted for by crowding of two proteins in the same bilayer.

BLTP1 forms a complex with the ER membrane proteins LTAP1 and TMEM170A^60,64^. An AlphaFold^73^ model of FIT2 with BLTP1(1–1500) placed the two chains at no confident interface (pTM = 0.49, ipTM = 0.19; Extended Data Fig. 6e), suggesting that FIT2 is not a constitutive subunit of the BLTP1 complex. This behavior is characteristic of a transient, lipid-context-dependent interaction that structure-prediction methods trained on stable complexes would fail to capture.

If the module is conditional rather than constitutive, its assembly should track the demand for LD biogenesis. Consistent with this, more endogenous FIT2 co-precipitated with full-length BLTP1–GFP after OA loading, at both 2 h and 15 h, whereas FIT2 levels in the input were unchanged (Fig. 3f). BLTP1(1–67)–GFP showed the same OA-dependent increase (Extended Data Fig. 6f), indicating that the N-terminal transmembrane segment suffices for this regulation. BLTP1 and FIT2 are therefore brought together on demand, when the cell is building LDs, rather than existing only as a preformed complex.

The *FITM2* missense variant p.Gly232Arg has been reported in the recessive deafness–dystonia syndrome associated with biallelic *FITM2* loss^48,49^. In cells, BLTP1-GFP formed discrete puncta that overlapped with mCh–FIT2 WT but not with mCh–FIT2-G232R (Fig. 3g,h). Importantly, GFP-Trap capture of BLTP1(1–67)–GFP efficiently pelleted Halo–FIT2 WT but only a trace of Halo–FIT2 G232R, although both were expressed at similar levels (Fig. 3i). Together, these experiments define a transmembrane BLTP1–FIT2 interface that is mediated by the TM region of BLTP1, is engaged on demand during LD biogenesis, and is strongly weakened by a single disease-associated substitution in FIT2.

### Both the FIT2 interface and the lipid-transfer groove are required for BLTP1 function

To ask whether the BLTP1–FIT2 interaction is essential for BLTP1 function in LD biology, we expressed GFP-tagged BLTP1 full-length and fragments in BLTP1-depleted cells and scored LD number and diameter (Fig. 4a–i).

**Fig. 4.**
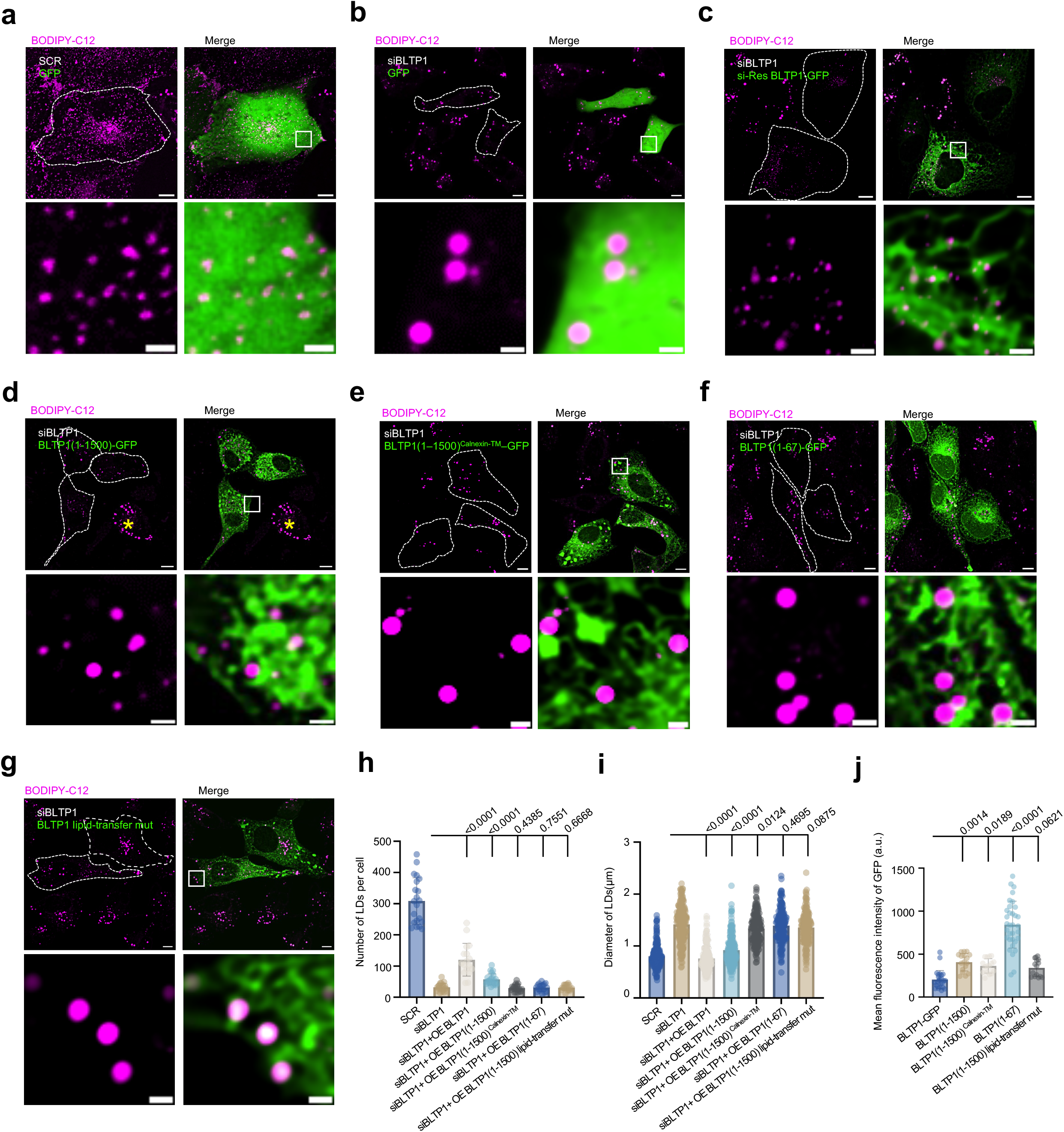
The N-terminal anchor and the lipid-transfer groove are both required for BLTP1 function in LD biology. **a–g**, Representative BODIPY images of oleate-loaded U2OS cells expressing: **a**, SCR + GFP vector; **b**, siBLTP1 + GFP vector; **c**, siBLTP1 + siRNA-resistant BLTP1–GFP; **d**, siBLTP1 + BLTP1(1–1500)–GFP; **e**, siBLTP1 + BLTP1(1–1500)^Calnexin-TM^–GFP; **f**, siBLTP1 + siRNA-resistant BLTP1(1–67)–GFP; **g**, siBLTP1 + BLTP1(1–1500)–GFP lipid-transfer mutant. **h**, LD number per cell from (a-g): SCR (21 cells); siBLTP1 (21 cells); siBLTP1+ full-length BLTP1-GFP (17 cells); siBLTP1+BLTP1(1–1500)-GFP (22 cells); siBLTP1+ BLTP1(1–1500)^Calnexin-TM^–GFP (20 cells); siBLTP1+BLTP1(1–67)-GFP (18 cells) and siBLTP1+BLTP1(1–1500) -GFP lipid-transfer mut (17 cells) from at least three independent experiments. Mean ± SD. Ordinary one-way ANOVA with Tukey’s multiple comparisons test. **i**, LD diameter in cells from (**a-g**) as (**h**). Mean ± SD. Ordinary one-way ANOVA with Tukey’s multiple-comparisons test. **j**, Mean GFP fluorescence intensity of cells expressing the indicated constructs, measured by flow cytometry. Mean ± SD. Ordinary one-way ANOVA with Tukey’s multiple-comparisons test. Scale bars: 10 μm in whole-cell image and 2 μm in insets.

All rescue experiments were scored against an internal control, comparing plasmid-transfected cells with non-transfected cells within the same imaging field. Control cells expressing GFP alone showed the normal LD population, and importantly, introduction of siRNA-resistant BLTP1-GFP, but not GFP alone, significantly rescued the LD phenotype (Fig. 4a–c,h,i). BLTP1(1–1500)-GFP, which carries the N-terminal transmembrane anchor together with the first ∼1,430 residues of the RBG rod, partially restored the defect (Fig. 4d,h,i), suggesting that the C-terminal region, including the region that binds the outer-mitochondrial-membrane receptor FKBP8^67^, is also required for BLTP1 function in LD biology. In addition, we replaced the N-terminal transmembrane segment of BLTP1(1–1500) with that of Calnexin, which anchors in the ER but does not efficiently engage FIT2 (Extended Data Fig. 7a,b), and expressed the resulting BLTP1(1–1500)^Calnexin-TM^–GFP chimera in BLTP1-depleted cells. Unlike BLTP1(1–1500)–GFP, this chimera failed to restore LD number and only slightly reduced LD diameter (Fig. 4e,h,i), suggesting that the interaction with FIT2 is required for BLTP1 function in LD biology. However, the TM anchor alone was insufficient, as BLTP1(1–67)–GFP rescued neither parameter (Fig. 4f,h,i).

We next asked whether lipid transfer itself is required for BLTP1 function. Following the strategy used to abolish lipid transfer by ATG2^57^, we introduced 37 substitutions of groove-lining hydrophobic residues into BLTP1(1–1500) (A382Q through F490Q; Extended Data Fig. 7c). This groove mutant failed to rescue either parameter: LD number and diameter were indistinguishable from those of cells given no rescue construct (*P* = 0.67 and 0.088; Fig. 4g–i). An intact groove is therefore indispensable for BLTP1 function. These failures were not due to insufficient expression. By flow cytometry, every construct that failed to rescue — the Calnexin-anchored chimera, BLTP1(1–67)–GFP and the groove mutant–was expressed at least as highly as full-length BLTP1–GFP, which rescued both parameters, with BLTP1(1–67)–GFP expressed highest (Fig. 4j). Together, both the FIT2-binding anchor and an intact groove are required for BLTP1 to support LD biogenesis.

### BLTP1 depletion reduces PA abundance and remodels the LD-associated lipidome

To determine whether BLTP1 depletion alters the lipid environment associated with LDs, we performed targeted LC–MS/MS on isolated LD fractions from control and BLTP1-depleted U2OS cells loaded with 200 µM OA. Five independent transfections were analyzed per condition, and lipid abundances were normalized to LD-fraction protein. The LD fractions contained the LD coat protein PLIN2 but no detectable plasma-membrane (ATP1A1), Golgi (GM130), lysosomal (LAMP1), mitochondrial (VDAC), ER (VAPB) or cytosolic (GAPDH) marker, all of which were readily detected in total cell lysates (Fig. 5a). We quantified 14 polar-lipid classes and free fatty acids in these fractions by targeted normal-phase LC–MS/MS against class-specific internal standards.

**Fig. 5.**
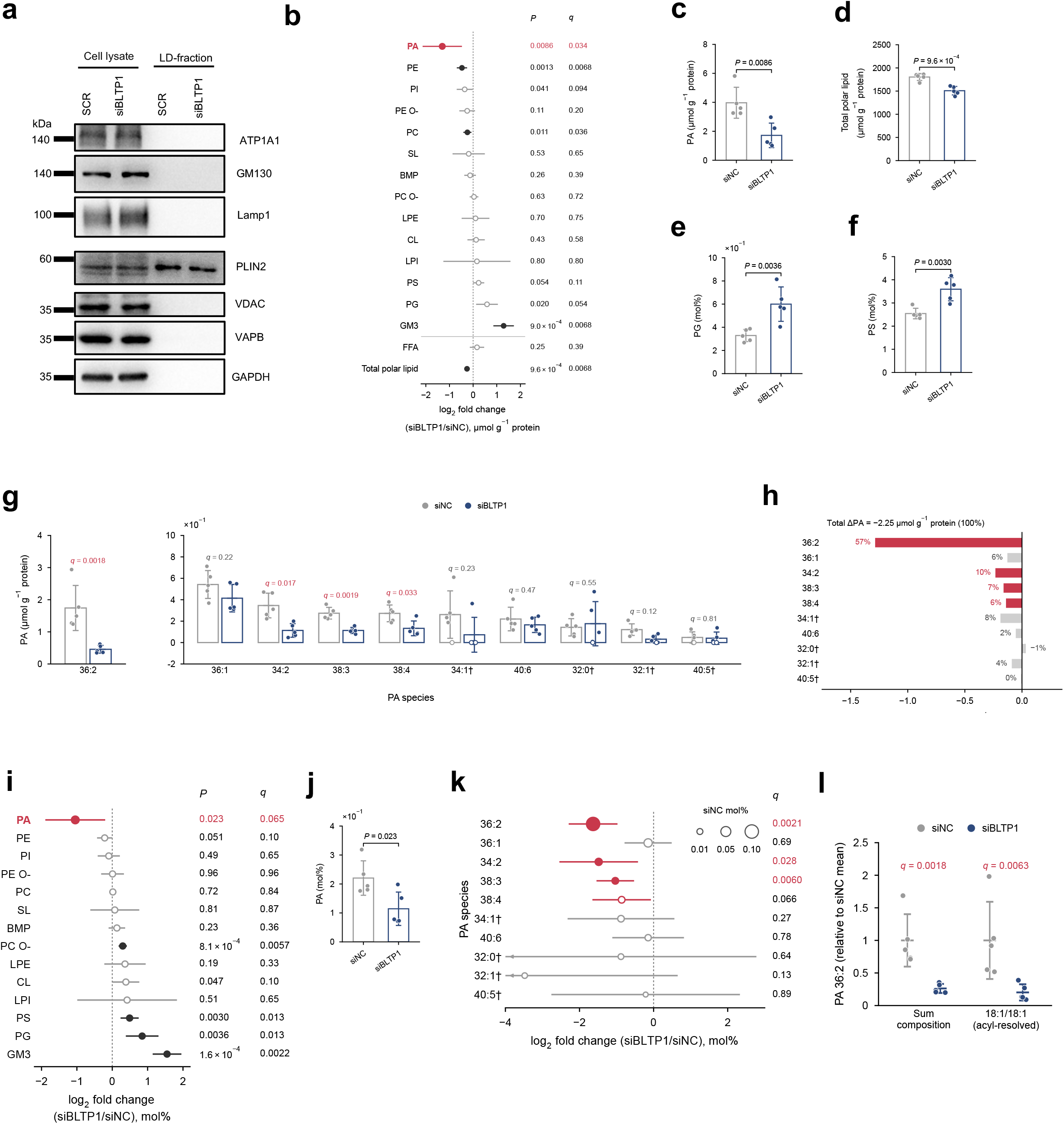
BLTP1 depletion selectively reduces PA in purified lipid droplets. **a**, Representative immunoblots of total cell lysates and floated LD fractions from U2OS cells transfected with SCR or siBLTP1 and loaded with 200 µM OA for 16 h from three independent experiments. The markers are PLIN2 (LD), ATP1A1 (plasma membrane), GM130 (Golgi), LAMP1 (lysosome), VDAC (mitochondria), VAPB (ER) and GAPDH (cytosol). **b**, Class-level change in LD-fraction lipids normalized to fraction protein. Points show log2 ratios of geometric means (siBLTP1/SCR) and lines 95% confidence intervals; PA is shown in red. Filled symbols denote *q* < 0.05 and open symbols *q* ≥ 0.05 (BH across the 16 features shown). Free fatty acids (FFA) and summed polar lipid are shown below the line. **c**,**d**, PA (**c**) and total polar lipid (**d**) per unit fraction protein. **e**,f, PG (**e**) and PS (**f**) as mol% of total measured polar lipid. **g**, PA sum-composition species per unit fraction protein, with PA 36:2 on a separate axis. Open dots mark values below detection. *q*, BH across 199 polar-lipid species. **h**, Contribution of each PA species to the absolute loss of PA, as the difference of group means. Species differences sum to the change in total PA. Red, species with *q* < 0.05 in **g**. **i**, As in **b**, with each class expressed as mol% of total measured polar lipid (BH across 14 classes). PA was the prespecified comparison. **j**, PA as mol% of polar lipid. **k**, Change in PA species expressed as mol%, ordered by abundance in SCR. Symbol area is proportional to SCR mol%, and lines show 95% confidence intervals. Red marks *P* < 0.05 and filled symbols *q* < 0.05 (BH across 199 polar-lipid species). **l**, PA 36:2 measured by its sum-composition transition and by its acyl-resolved 18:1/18:1 transition, each normalized to the SCR mean. Horizontal marks and whiskers show the Mean ± SD *q* values are BH-adjusted across 199 polar-lipid species (sum composition) or across the nine acyl-resolved PA transitions (18:1/18:1). Both channels measure the same samples. In **c**–**f** and **j**, bars show the mean, error bars the SD and dots individual LD preparations; **g** is displayed in the same way. n = 5 LD preparations per group, each from an independent transfection. *P* values are from two-sided Welch’s *t*-tests on log2-transformed values. † marks species with at least one value below detection, imputed as half the species minimum for statistics. PE O- and PC O-, ether-linked PE and PC; SL, sulfatide; LPE and LPI, lyso-PE and lyso-PI; BMP, bis(monoacylglycero)phosphate.0

Normalized to fraction protein, PA showed the largest proportional decrease of any lipid class, falling from 4.0 ± 1.1 to 1.7 ± 0.8 µmol g^−1^ protein (mean ± SD; 0.43-fold; *P* = 0.0086, *q* = 0.034; Fig. 5b,c). PE and PC decreased less (0.72-fold and 0.85-fold; *q* = 0.0068 and 0.036), and total polar lipid fell by 16%, from 1,803 ± 75 to 1,508 ± 92 µmol g^−1^ protein (*P* = 9.6 × 10^−4^; Fig. 5b,d). Among the anionic phospholipids, the loss was specific to PA. As a proportion of polar lipid, PG and PS increased 1.8-fold and 1.4-fold (*q* = 0.013 for both; Fig. 5e,f).

At the species level, four PA species decreased significantly (Fig. 5g): PA 36:2 (0.26-fold; *q* = 0.0018), PA 38:3 (0.41-fold; *q* = 0.0019), PA 34:2 (0.33-fold; *q* = 0.017) and PA 38:4 (0.49-fold; *q* = 0.033). The remaining six species, including the abundant PA 36:1 and PA 40:6, did not change significantly. PA 36:2, the most abundant PA species in control LDs, alone accounted for 57% of the total PA loss of 2.25 µmol g^−1^ protein (Fig. 5h).

Normalization to fraction protein assumes that the amount of co-purifying protein per unit of LD monolayer is the same in both conditions. This may not hold for LDs of different size and coat composition. We therefore also expressed each class as mol% of total measured polar lipid, a measure of monolayer composition that does not depend on this denominator. PA remained the most strongly reduced class, falling from 0.22 ± 0.06 to 0.11 ± 0.06 mol% (*P* = 0.023 for this prespecified comparison; *q* = 0.065 across 14 classes; Fig. 5i,j), whereas PC was unchanged. PA 36:2 (0.32-fold; *q* = 0.0021), PA 34:2 (*q* = 0.028) and PA 38:3 (*q* = 0.0060) again decreased (Fig. 5k), so these three species changed with the same direction and significance under both normalizations.

The acyl-chain-resolved transition recorded for PA 36:2 corresponds to 18:1/18:1, consistent with de novo PA assembled from acyl-CoA through the GPAT/AGPAT route using the OA supply^24,25^. This transition decreased 0.20-fold (*q* = 0.0063), in line with the sum-composition measurement (0.26-fold; *q* = 0.0018; Fig. 5l). Selective loss of this species therefore points to the actively synthesized, biogenesis-associated PA pool rather than to the pre-existing structural PA of the LD monolayer.

BLTP1 depletion also remodeled the LD-fraction lipidome more broadly. The two conditions separated completely along the first principal component (50.3% of variance; Extended Data Fig. 7d), and 105 of 199 species differed at *q* < 0.05 (73 increased and 32 decreased; Extended Data Fig. 7e). Notably, PA 36:2 was the most strongly reduced species that passed the 5% false-discovery threshold.

The targeted method measures PA, PE, PE O-, PI and PS through two separate sets of transitions: sum-composition species and acyl-chain-resolved species. Class sums from the two channels agreed for PA (0.40-fold and 0.29-fold) and PE (0.72-fold and 0.69-fold) but diverged for PI, PS and PE O-(Extended Data Fig. 7f). The PA decrease is therefore independent of the measurement channel, whereas the small class-level changes in PI, PS and PE O-depend on the channel and are not interpreted further.

We next asked whether BLTP1 depletion also lowers PA in the bulk ER. ER fractions were isolated from SCR and siBLTP1-treated U2OS cells loaded with OA. These fractions were enriched for VAPB and contained only traces of GM130, LAMP1 and GAPDH, with no detectable pan-Cadherin, PLIN2 or VDAC (Extended Data Fig. 8a). In contrast to LDs, ER PA was unchanged whether normalized to fraction protein (4.8 ± 0.6 versus 5.1 ± 0.5 µmol g^−1^ protein; *P* = 0.38; Extended Data Fig. 8b,c) or expressed as mol% of polar lipid (0.69 ± 0.09 versus 0.75 ± 0.10 mol%; *P* = 0.31; Extended Data Fig. 8j,k). No ER PA species changed at *q* < 0.05 (Extended Data Fig. 8o). Total polar lipid per unit protein, the major classes PC, PE, PI and PS, and CL were likewise unchanged (Extended Data Fig. 8b,d,f,g,j,m,n).

Changes in the ER were confined to minor classes. PG and GM3 increased and sulfatide decreased under both normalizations (per protein: 1.44-fold, 1.89-fold and 0.44-fold; *q* = 0.023, 0.021 and 0.0056; Extended Data Fig. 8b,e,j,l), BMP rose modestly and ether-linked PC fell slightly in mol% (1.20-fold and 0.88-fold; *q* = 0.033 and 0.024), but neither changed per protein (Extended Data Fig. 8b,h,j). Because PG and GM3 rose in both the LD and ER fractions, those changes appear not to be restricted to LD fractions; the PA decrease, by contrast, was specific to the LD fraction. Despite unchanged class totals, 90 of 208 ER species differed at *q* < 0.05 (Extended Data Fig. 8i,p), indicating acyl-chain remodeling within the major ER phospholipid classes.

Together, these data show that BLTP1 is required to maintain PA in LD-associated membranes but not in the bulk ER, and that among the anionic phospholipids this requirement is specific to PA. The loss is concentrated in PA 36:2, the species consistent with PA made from the supplied OA.

### BLTP1 is required for adipogenic lipid droplet accumulation

LD biogenesis factors are most stringently required in cells specialized for lipid storage. Seipin loss, for example, causes generalized lipodystrophy in humans^11^, and FIT2 is required for normal adipose triglyceride storage in mice^44^. We therefore asked whether BLTP1 is required for adipogenic LD accumulation. Human adipose-derived stem cells (ADSCs) were transfected with control siRNA or siRNAs targeting BLTP1, FIT2, FTO or Seipin/BSCL2, induced to differentiate, and imaged by BODIPY and DIC at days 0, 4, 8, 12 and 16 (Fig. 6a–e). Control cells accumulated large, BODIPY-bright LDs over the time course. In contrast, all four knockdowns reduced the fraction of BODIPY-positive, differentiated cells throughout ADSC differentiation (Fig. 6f). Transcript levels of the adipogenic markers aP2, PPARγ, adipsin, C/EBPα, LPL and leptin were also generally lower in the knockdown cells (Fig. 6g–l). Knockdown was confirmed by RT–qPCR (Fig. 6m). These experiments suggest that BLTP1 and FIT2 are each required for adipogenic LD accumulation.

**Fig. 6.**
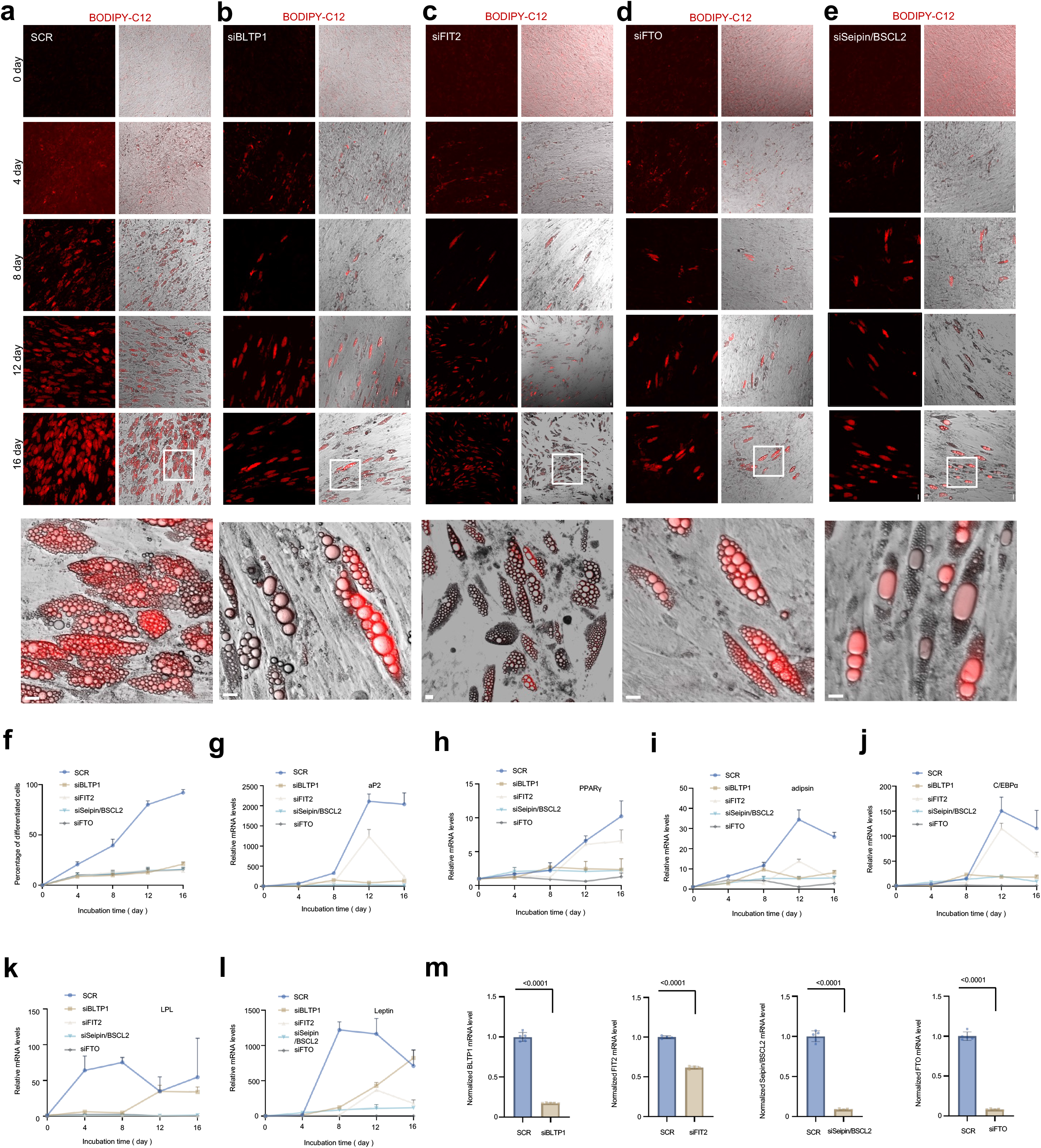
BLTP1 supports adipogenic LD accumulation. **a–e**, Representative BODIPY and DIC images of human adipose-derived stem cells at days 0, 4, 8, 12 and 16 of adipogenic differentiation, transfected with SCR (**a**), siBLTP1 (**b**), siFIT2 (**c**), siFTO (**d**) or siSeipin/BSCL2 (**e**). **f**, Fraction of differentiated (BODIPY-positive) cells during differentiation (days 0–16). Mean ± SD. **g–l**, RT-qPCR of adipogenic markers aP2 (**g**), PPARγ (**h**), adipsin (**i**), C/EBPα (**j**), LPL (**k**) and leptin (**l**). Mean ± SD. **m,** qPCR showing the efficiency of suppression of BLTP1, FIT2, Seipin/BSCL2 and FTO in cells as in (**a–e**) from at least three independent experiments. Mean ± SD. Two-tailed unpaired Student’s t-tests. Scale bars: 10 μm in whole-cell image and 2 μm in insets.

## Discussion

Models of LD biogenesis have focused on the protein machinery that concentrates neutral lipid at defined ER sites. Our results add the lipid supply line that serves those sites, complementing the recent finding that ATG2A delivers diacylglycerol to recruit DGAT2 to growing LDs^74^. BLTP1 is required for cells to build a normal number of LDs and to keep the LDs they build from overgrowing; it engages the emergence factor FIT2 through its N-terminal TM segment, an interface that is strengthened when LD biogenesis is induced and strongly weakened by a disease-associated FIT2 substitution. Both the FIT2 interface and an intact transfer groove are required for BLTP1 to function. Suppressing FIT2 relieves the overgrowth caused by BLTP1 loss, consistent with the two proteins acting in sequence rather than in parallel. BLTP1 loss most strongly drains PA from purified LD fractions while leaving ER PA and the major ER phospholipid classes unchanged. We propose that BLTP1-mediated local PA supply to LD formation sites on the ER determines how efficiently LDs nucleate and how large they grow (working model; Extended Data Fig. 9).

That two interacting proteins give opposite LD phenotypes is what an ordered pathway predicts. FIT2 depletion gave few, small LDs whereas BLTP1 depletion gave few, large ones (Fig. 1e–g). FIT2 acts first, letting neutral lipid into the nascent LDs^37,42^, so losing it starves LDs of the material they need to grow. BLTP1 acts on the LDs that FIT2 has allowed to form, sustaining the PA that shapes their surface^3–5,23^; when only BLTP1 is missing, lipid continues to arrive but reaches too few droplets, which therefore overgrow. Removing both blocks the influx as well, which is why the double depletion retains the low LD number of BLTP1 loss but loses most of the size increase. The neutral-lipid measurements follow the same order. FIT2 depletion lowered cellular triacylglycerol, whereas BLTP1 depletion raised it while reducing droplet number (Fig. 2h,i). More storage lipid in fewer, larger LDs is the expected outcome when nucleation fails but delivery does not^3,21,26^.

The lipidomic result argues that this handling is site-restricted. Purified LD fractions from BLTP1-depleted cells lost PA specifically, whereas ER fractions from the same cell background showed no change in PA or in the major phospholipid classes (Fig. 5 and Extended Data Fig. 8). A protein that supplied phospholipid to the ER as a bulk pool^61,62^ would not be expected to leave the bulk ER composition intact, so BLTP1 activity relevant to LD formation is confined to a subdomain that fractionates with LDs. PA is the most mechanistically loaded of these species: it is anionic, has negative intrinsic curvature, lowers oil–water interfacial tension, and both its abundance and its asymmetry across the leaflets influence whether a lens forms and in which direction it buds^4–7,16^; it is the substrate consumed by Lipin/PAP en route to diacylglycerol and triglyceride^24,25^; and PA-enriched ER–mitochondria subdomains recruit Seipin through ORP5/ORP8^30^. PA excess drives supersized LDs in yeast^26^, whereas in this study PA is depleted and LDs are larger, so the size phenotype is unlikely to reflect PA-driven fusogenicity and more likely reflects loss of a monolayer surfactant environment that normally restrains coalescence and ripening^3,21,23^. Distinguishing these requires measuring PA where it acts rather than in a fraction.

It is still unclear where this BLTP1-FIT2 module sits relative to Seipin–LDAF1. BLTP1–FIT2 may work inside the Seipin-marked assembly site, feeding PA to the same nucleation event the Seipin cage catalyses^15,17,19^. It may instead mark a separate, FIT2-dependent subdomain^41^. Or it may act earlier, setting the lipid environment that decides whether a Seipin-marked lens ever matures^10,21^. Each predicts something different for Seipin and LDAF1 puncta density, dwell time and maturation rate after BLTP1 loss, which is how we would tell them apart.

Where does this PA come from? Mitochondria-associated ER membranes (MAM) are an established platform for LD assembly. ORP5 and ORP8 recruit Seipin to PA-enriched MAM subdomains^30^. ORP5 also occupies ER–LD contacts^29^. MIGA2 tethers mitochondria, ER and LDs to support lipogenesis^32^, and the bridge-like transporter VPS13A is found at ER–mitochondria–LD contacts^33^. A recent ORP5 preprint reports mitochondria-to-ER/MAM PA transfer at mitochondria–ER–LD junctions^75^. A mitochondrial PA pool delivered to the ER at MAM is therefore a plausible source for the PA that LD biogenesis consumes, and we propose it as a route worth testing directly.

BLTP1 is well suited to participate in such a route. It is a rod long enough to span a contact site, with an ordered lipid-filled groove and an associated ER scramblase^52,53,60,64^, and it carries spatially separate adaptors at its two ends. FIT2 engages the N-terminal TM segment characterized here, whereas the outer-mitochondrial-membrane protein FKBP8 binds the C-terminal region in a companion study^67^. A single bridge with two receptors can be positioned at two different interfaces, and the present work concerns only the N-terminal, FIT2-dependent, ER–LD arm. Whether the mitochondrial and ER–LD arms of BLTP1 are connected, and in which direction lipid moves between them, is the obvious next question warranting further investigation (working model; Extended Data Fig. 9).

The BLTP1–FIT2 interface also links two clinically distinct disorders. Biallelic BLTP1/KIAA1109 variants cause Alkuraya–Kučinskas syndrome, characterized by severe neurodevelopmental impairment, hypotonia and arthrogryposis and frequently lethal before or shortly after birth^65^; Bltp1 loss is perinatally lethal with neuromuscular defects in mice^66^, and the fly orthologue Tweek is required for phosphoinositide-dependent membrane traffic^63^. Biallelic *FITM2* variants cause a postnatal, progressive deafness–dystonia syndrome with sensorineural hearing loss, generalized dystonia, motor regression and ichthyosis^48,49^, and FIT2 loss disrupts ER homeostasis in intestine, liver and β cells^45–47^. Their partial convergence on developmental and motor dysfunction suggests shared tissue vulnerability to disturbed organelle lipid homeostasis. The ability of FIT2-G232R to weaken BLTP1 association provides a tractable hypothesis for testing in patient-derived cells.

## Methods

### Cell culture, transfection and RNAi

Human U2OS (osteosarcoma), HeLa (cervical carcinoma), HepG2 and Huh7 (hepatocellular carcinoma) and HEK293 cells were obtained from ATCC and maintained in DMEM (Gibco) supplemented with 10% fetal bovine serum (Gibco) at 37 °C in 5% CO2. Cell lines were authenticated by the supplier and confirmed free of mycoplasma by PCR at the beginning and end of the study. For plasmid transfection, cells were seeded at 4 × 10^5^ per well of a six-well plate ∼16 h before transfection and transfected in Opti-MEM (Invitrogen) with polyethyleneimine (4 µL per µg DNA) for 5 h, then trypsinized and replated onto 35-mm glass-bottom dishes at ∼3.5 × 10^5^ cells per dish. Cells were imaged 16–24 h after transfection in imaging medium (phenol red-free DMEM, 10% FBS, 20 mM HEPES, no antibiotics). Per 35-mm dish we used 1,000 ng of BLTP1-GFP or BLTP1 truncation constructs and 300 ng of mCherry-FIT2, Halo-FIT2, mCherry-LiveDrop, Halo–Sec61β or other marker constructs, individually or combined.

For RNAi, cells were plated at 30–40% confluence and transfected with 50 ng siRNA and 2 µL Lipofectamine RNAiMAX (Invitrogen) per 35-mm dish; a second identical round was performed 48 h later and cells were analyzed 24 h after the second transfection. siRNA sequences (Ribobio) were: scrambled 5′-CGUUAAUCGCGUAUAAUACGCGUAT-3′; BLTP1 #1 5′-GCATGAAGTTGACCATTAA-3′; BLTP1 #3 5′-CCAGAACTCTGCCATTCAA-3′; FIT2 #1 5′-UCAUUGCCCUCACCAACUATT-3′; FIT2 #2 5′-CCUCAUGAUUGUUGAAGAGTT-3′; *Seipin/BSCL2* #1 5’-GCUCCUUCUACUAUUCCUATT-3’; *Seipin/BSCL2* #2 5’-GCAUGUUCUUGGUCACCAUTT-3’; *Seipin/BSCL2* #3 5’-CCUCCUGCUAUUUGGCUUUTT-3’; FTO #1 5’-GCUGAAAUAUCCUAAACUATT-3’. For double knockdown, BLTP1 and FIT2 siRNAs were co-transfected at 50 ng each, and single-knockdown samples were balanced with scrambled siRNA to equal total siRNA. Knockdown of BLTP1 was quantified by RT–qPCR (below). FIT2 knockdown efficiency was quantified by Western blots or RT–qPCR in this study.

### Plasmids and cloning

Human BLTP1 (NM_001384125.1) was synthesized (Synbio Technologies); human FIT2 (FITM2, NM_001080472.4) was amplified from HEK293 cDNA. Full-length BLTP1 was cloned into mEGFP-N1 (Addgene 54767) between SalI and SacII. FIT2 was cloned into mCherry-C1, Halo-C1 and 3×Flag vectors between HindIII and BamHI. BLTP1 truncations (1–67, 68–1500, 1–1500, 1501–2610, 2611– 3592, 3593–5093) were cloned into GFP-N1, and FIT2 truncations (1–141, 87–262, 142–262) into Halo-C1, between HindIII and BamHI. FIT2 point mutants G232R, corresponding to the FITM2 c.694G>A allele reported in Siddiqi syndrome^49^ were generated by overlap-extension PCR. The BLTP1(1–1500)-GFP groove mutant carrying 37 substitutions of groove-lining hydrophobic residues (A382Q through F490Q) and the BLTP1(1–1500)^Calnexin-TM^-GFP chimera, in which residues 1–67 were replaced by the calnexin transmembrane segment, were generated by gene synthesis and verified by full-insert Sanger sequencing. mCherry-LiveDrop and LiveDrop-GFP encode the *Drosophila* GPAT4 amphipathic hairpin^8,10^. Halo–Sec61β, GFP–Sec61β, mCherry-RTN4a, Halo-ATL3, ACSL3-Halo, LDAF1-mCherry, BSCL2-mCherry, mRuby-SigmaR1, BLTP1(1–67)-mClover, FIT2-mRuby and RFP-GFP-PLIN2 were used as markers or controls. All primers are listed in Supplementary Table 1.

**Supplementary Table 1.**
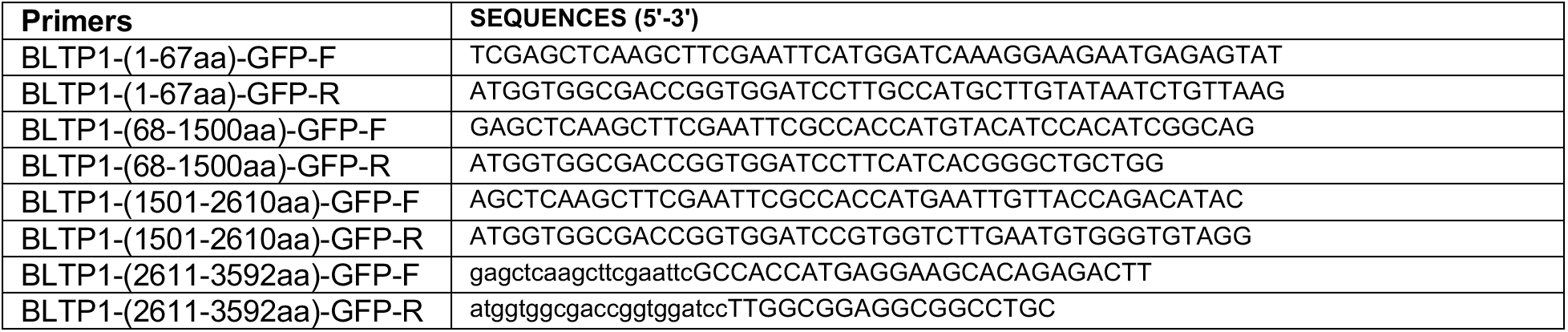
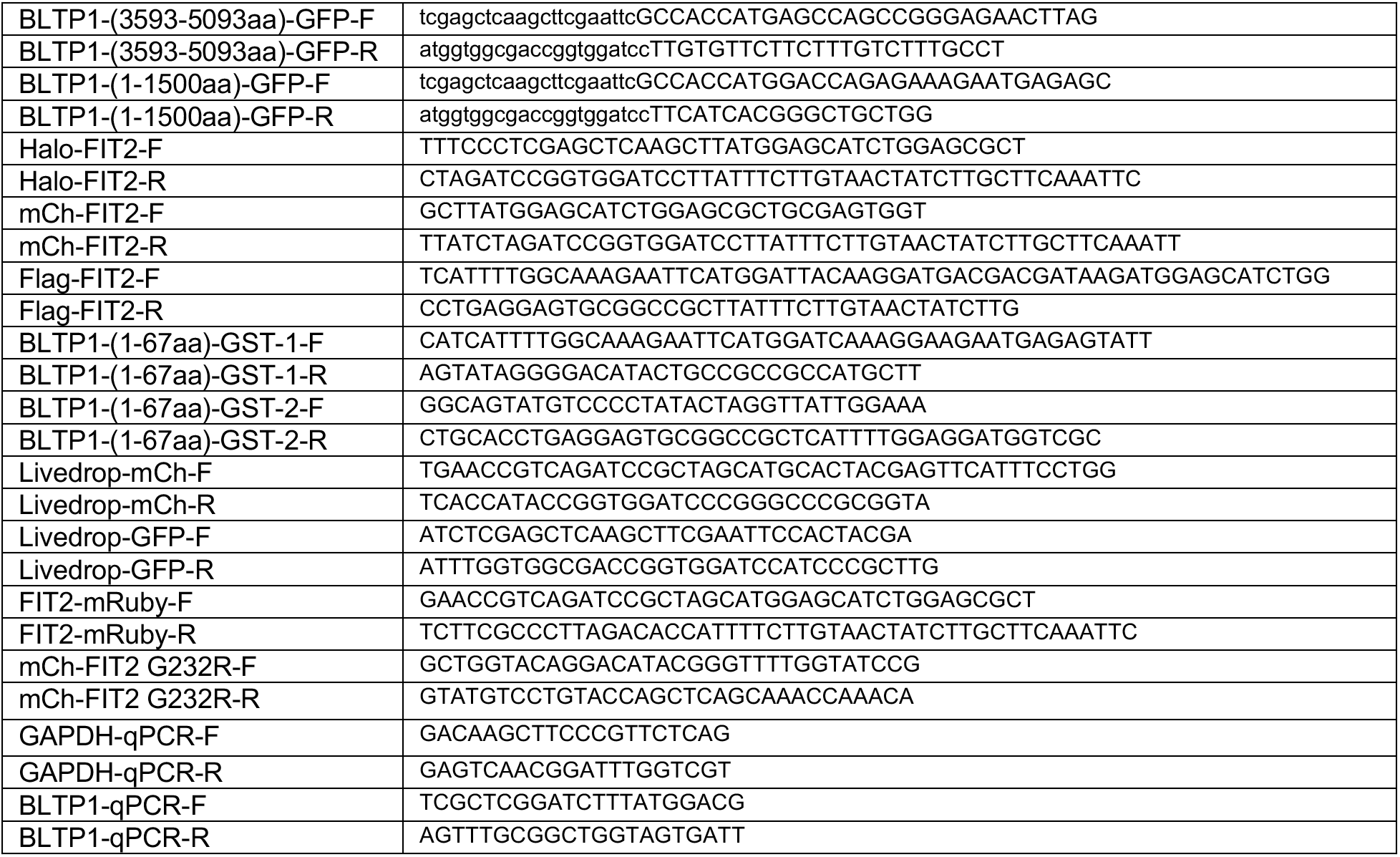
Primers used in this study.

### CRISPR–Cas9 knock-in line

The endogenous BLTP1-3×Flag knock-in U2OS line was generated by electroporation of recombinant Cas9 protein, sgRNAs (5′-TAAGACTTCCTCTAGAACTG-3′ and 5′-TCTAGAGGAAGTCTTAATGG-3′) and a single-stranded homology-directed repair oligonucleotide encoding a C-terminal 3×Flag tag. Monoclonal lines were expanded and the in-frame insertion confirmed by Sanger sequencing of genomic PCR amplicons and by anti-Flag dot blotting.

### Human ADSC differentiation

Human adipose-derived stem cells (ADSCs) were grown to confluence in 24-well plates and induced with an adipogenic cocktail (Procell PD-006) containing 3-isobutyl-1-methylxanthine, dexamethasone, human insulin and rosiglitazone for 3 d, then switched to maintenance medium (insulin, glutamine, penicillin–streptomycin) for 1 d; the two media were alternated until day 16.

siRNAs against BLTP1, FIT2, FTO or Seipin/BSCL2 were introduced by RNAiMAX transfection before induction and re-transfected every 96 h. Cells were imaged by confocal and DIC microscopy at days 0, 4, 8, 12 and 16 after BODIPY-C12 labeling, and adipogenic markers (FABP4/aP2, PPARG, CFD/adipsin, CEBPA, LPL, LEP) were measured by RT–qPCR. For the differentiation index in Fig. 6f, cells were scored as differentiated or undifferentiated from paired DIC and BODIPY images. Within each large field-of-view image, the total number of cells and the number of cells positive for BODIPY staining were counted. The percentage of BODIPY–positive cells was then calculated relative to the total cell population. Data are expressed as the percentage of BODIPY–positive cells among total cells.

### RT–qPCR

Total RNA was extracted with TRIzol (Thermo Fisher), reverse-transcribed with ReverTra Ace (TOYOBO, TRT-101) and amplified with qPCR Mix (TOYOBO, QPS-201) on a Bio-Rad CFX system. Relative expression was calculated by the 2^−ΔΔCt^ method with GAPDH as the reference gene. Primers are listed in Supplementary Table 1. BLTP1 knockdown was validated by RT–qPCR in the same biological replicates used for imaging.

### Immunoblotting and dot blotting

For SDS–PAGE, cells were lysed directly in SDS sample buffer and heated to 95 °C for 10 min, resolved on 8–12% gels, transferred to a nitrocellulose membrane (Millipore), blocked in 5% non-fat milk for 1 h at room temperature, incubated with primary antibodies overnight at 4 °C and with HRP-conjugated secondaries (1:10,000, Sangon D110087/D110058) for 1 h, and developed with ECL (Beyotime P0018M-2). Antibodies were: GFP (ABclonal AE011), mCherry (ABclonal AE002), Halo (Promega G928A), FIT2 (Cell Signaling 62622), RTN4 (ABclonal A1752), ATL3 (Solarbio K008635P), Flag (Proteintech 66008), PLIN2 (A6276, ABclonal), VAPB (Proteintech 14477-1-AP), LAMP1 (sc-20011, Santa Cruz), ATP1A1 (Proteintech 83191-6-RR), VDAC (A0810 Abclonal), GM130 (M179-3, MBL), Tubulin (100109-MM05T, Sino Biological Inc.), BiP (Proteintech 66574-1-Ig), Actin (Proteintech 66009-1-Ig), Pan-cadherin (Proteintech 31313-1-AP), GST (Proteintech 66001-2-Ig) and GAPDH (AB9485, Abcam). All antibodies were used at 1:2000 for western blots. Endogenous BLTP1 has a predicted mass of ∼560 kDa and does not enter or resolve reproducibly in standard SDS–PAGE, so it cannot be quantified by conventional immunoblotting. BLTP1-3×Flag was therefore detected by dot blotting: cleared lysates were normalized by BCA assay, spotted (2 µL, 5–20 µg total protein) onto nitrocellulose, air-dried, blocked and probed with anti-Flag exactly as for immunoblots, with GAPDH spots as loading control.

### ER stress assays

Thapsigargin-treated and vehicle-treated cells were compared for BiP/HSPA5 levels by immunoblotting and for ER-whorl formation by confocal imaging of an ER marker (Extended Data Fig. 2). Thapsigargin was purchased from Abcam and dissolved in DMSO to prepare a 1.53 mM stock solution. Cells were treated with 2 μM thapsigargin for 3, 6, 9, or 12 h, with DMSO used as the vehicle control. ER whorls were defined morphologically as transformation of the ER into large ring-like structures.

### Autophagy, lysosome and LD-turnover assays

LC3B puncta, LysoTracker-positive compartments and their apposition to LDs, and acidification of the tandem RFP–GFP–PLIN2 reporter, were assessed by live-cell confocal imaging (Extended Data Fig. 3). For lysosome labeling, cells were loaded with LysoTracker™ Deep Red (cat. no. L12492) at a final concentration of 1 μM, prepared by dilution from a 1 mM stock solution, and incubated for 30 min at 37 °C in a 5% CO₂ incubator. For the operational definition of LD–lysosome or LD–LC3B contacts, spatial co-localization of the two markers was required, and contact was defined as overlap of the radii of the two markers. RFP–GFP–PLIN2 puncta were classified as acidified when they were positive for RFP signal and negative for GFP signal.

### Live-cell confocal microscopy

Cells on 35-mm glass-bottom dishes were imaged on a Zeiss LSM780 laser-scanning confocal microscope equipped with 405, 458, 488, 514, 561 and 633 nm lasers and a spectral GaAsP detector, using a 63×/1.4 NA Plan-Apochromat oil objective, in a stage-top chamber at 37 °C and 5% CO2. Excitation was 405 nm for BFP, 488 nm for GFP/mClover/BODIPY 493/503, 561 nm for BODIPY-C12 558/568, mCherry and mRuby, and 633 nm for Janelia Fluor 646. Acquisition settings were held constant across all conditions within an experiment, and control and knockdown dishes were imaged in interleaved order on the same day.

### Structured illumination microscopy

Super-resolution imaging of mCherry-LiveDrop, Halo-Sec61β and BODIPY (Fig. 2e) was performed by three-dimensional structured illumination microscopy on live cells maintained at 37 °C. Raw stacks were reconstructed with the manufacturer’s algorithm, and reconstruction quality was assessed before quantification; identical illumination, exposure and reconstruction parameters were applied to all conditions within an experiment.

### Fluorescence lifetime imaging microscopy

U2OS cells were co-transfected with BLTP1(1–67)-mClover (donor) and FIT2-mRuby or mRuby-SigmaR1 (acceptors), or with donor alone, and imaged 24 h later on a Leica Stellaris confocal system. Donor lifetimes were fitted with the TauSense module^72^; mean lifetimes were extracted per cell over whole-cell ER regions of interest and averaged per independent experiment.

### BODIPY-C12, LipidTox and HaloTag labeling

For LD labeling, cells were washed once in PBS and incubated in complete medium containing 1 µM BODIPY 558/568-C12 for 20 min at 37 °C, washed three times in PBS and returned to imaging medium^68^. Neutral lipid staining in fixed samples used HCS LipidTox or BODIPY 493/503 per the manufacturer’s instructions. HaloTag fusions^70^ were labeled with 5 nM Janelia Fluor 646 HaloTag ligand^71^ for 30 min, washed three times in complete medium, chased for a further 30 min and switched to imaging medium immediately before acquisition.

### Delipidation and oleate-induced LD formation

Cells were delipidated for 72 h in DMEM containing 5% lipoprotein-deficient serum (Sigma) in place of FBS, then LD formation was induced by adding BSA-conjugated oleic acid (OA) for 0, 15, 30 or 60 min, or for the times indicated. In the delipidation and re-loading experiments (Fig. 2), cells were subjected to a 24-h delipidation prior to siRNA knockdown. At 24 h after the initiation of delipidation, siRNA transfection was performed, and delipidation was continued for an additional 48 h. At the end of the 72-h delipidation period, 200 μM (final concentration) OA was added, and cells were subsequently treated with OA for the indicated durations. For all OA experiments presented in Figs. 1, 2, 4, 5 and Extended Data Figs. 1–3, BSA-conjugated oleic acid (OA) was prepared from a 10 mM OA stock solution, with an OA: BSA conjugation ratio of 1:1, and was used at a final working concentration of 200 μM. These conditions were applied consistently across all OA experiments.

### Transmission electron microscopy

Cells were fixed in 2.5% glutaraldehyde in 0.1 M phosphate buffer pH 7.4 for 2 h at room temperature, washed three times, scraped, pelleted at 956g, post-fixed in 1% OsO4 for 2–3 h at 4 °C, dehydrated through graded ethanol (50–100%) and infiltrated with acetone–epoxy (2:1, 1:1) and pure epoxy for 12 h each at 37 °C before embedding. Sections of 60–100 nm parallel to the monolayer were cut on a Leica EM UC7 and imaged on a Tecnai G2 20 TWIN (FEI) at 200 kV. LDs were identified morphologically, traced manually and their number per cell section and cross-sectional area measured in ImageJ; diameters were calculated from areas assuming circular profiles. Five cell sections per condition were analyzed by an operator blinded to treatment condition.

### Lipid droplet and ER membrane fractionation

LD fractions were purified by sucrose density-gradient flotation^76^ with modifications. Cells were washed, lysed on ice in hypotonic lysis medium (HLM: 20 mM Tris-HCl pH 7.4, 1 mM EDTA, protease inhibitors) with a Potter–Elvehjem homogenizer and centrifuged at 1,000g for 10 min at 4 °C. The post-nuclear supernatant plus floating fat layer was adjusted to 20% sucrose with HLM/60% sucrose, dispersed thoroughly, loaded at the bottom of an SW41Ti tube and overlaid sequentially with 5 mL HLM/5% sucrose and 5 ml HLM, then centrifuged at 28,000g for 30 min at 4 °C. The floating LD fraction was recovered, re-pelleted at 20,000g for 10 min and resuspended in HLM. ER-enriched membranes were prepared by Percoll-gradient subfractionation^77^ from separately prepared homogenates of the same cell background. Fraction purity was assessed by immunoblotting for PLIN2^69^ (LD), VAPB (ER), VDAC (mitochondria), GM130 (Golgi), LAMP1 (endolysosome), ATP1A1 (plasma membrane) and GAPDH (cytosol).

### Recombinant protein expression and purification

Purified Flag-FIT2 and BLTP1(1–67)-GST were used for the in vitro binding experiments in Fig. 3b. Flag-FIT2 and BLTP1(1–67)-GST were expressed in Expi293 cells (Invitrogen) according to the manufacturer’s instructions for 72 h. The harvested cells were resuspended in lysis buffer: 25 mM HEPES pH 7.5, 150 mM NaCl and 1% protease-inhibitor cocktail (Roche), and lysed by sonication on ice. The membrane was solubilized by the addition of 1% (w/v) DDM and 0.1% cholesteryl hemisuccinate Tris salt at 4 °C for 2 h. After centrifugation at 20,000 g for 1 h, the supernatant was collected and loaded into pre-equilibrated anti-Flag M2 affinity resin (Sigma). The resin was washed with washing buffer containing 25 mM HEPES pH 7.5, 150 mM NaCl, 0.01% DDM and 0.001% CHS, and eluted with washing buffer supplemented with 250 μg/mL 3 × FLAG peptide.

### GST pull-down and GFP-Trap binding assays

For GFP-Trap and Flag-Trap co-immunoprecipitation from cells, monolayers were lysed in 50 mM Tris-HCl pH 7.5, 150 mM NaCl, 0.5 mM EDTA, 0.5% Nonidet P40 substitute with protease inhibitors; lysates were cleared at 20,000g for 10 min at 4 °C and incubated with GFP-Trap agarose (ChromoTek) or anti-Flag resin for 4 h at 4 °C with gentle rocking. Beads were washed five times in 50 mM Tris-HCl pH 7.5, 150 mM NaCl, 0.5 mM EDTA with protease inhibitors, eluted in SDS sample buffer and analyzed by immunoblotting; 5% input was loaded unless otherwise indicated. Endogenous co-immunoprecipitation (Fig. 3a) used the BLTP1-3×Flag knock-in U2OS line, with cells expressing the empty 3×Flag-N1 vector as negative control. For the GST pull-down shown in Fig. 3b, purified GST or BLTP1(1–67)-GST was incubated with GST beads for 2 h. The beads were then washed five times with HNM buffer (20 mM HEPES, pH 7.4, 0.1 M NaCl, 5 mM MgCl₂, 1 mM DTT, and 0.2% NP-40). Purified Flag-FIT2 was subsequently added, and the mixture was incubated for another 2 h, followed by five additional washes with HNM buffer. For the in vitro binding assays (Fig. 3c), HEK293 cells transiently transfected with BLTP1(1-67)-GFP were lysed in high-salt lysis buffer (RIPA buffer containing 500 mM NaCl, protease inhibitors and PMSF). GFP-Trap beads were used to pellet BLTP1(1-67)-GFP from cell lysates, followed by washing with high-salt lysis buffer for 10 times. The GFP beads were incubated with purified Flag-FIT2 2h at 4°C, respectively, followed by washing beads with freshly prepared HNM buffer. GST or GFP beads were resuspended in 100 μL 2 x SDS-sampling buffer. Re-suspended beads were boiled for 10 min at 95°C to dissociate protein complexes from beads. Western blotting was performed using anti-GFP, GST or Flag antibodies. The Coomassie staining was performed for purified GST, BLTP1(1–67)-GST or Flag-FIT2.

### Targeted LC–MS/MS lipidomics

Lipids were extracted from the ER and LD samples using a modified version of the Bligh and Dyer method as described previously^78^. Briefly, the ER and LD samples were homogenized in 750 µL of chloroform: methanol: MilliQ H_2_O (3:6:1) (v/v/v). The homogenate was then incubated at 1500 rpm for 1h at 4 ℃. At the end of the incubation, 350 µL of deionized water and 250 µL of chloroform were added to induce phase separation. The samples were then centrifuged and the lower organic phase containing lipids was extracted into a clean tube. Lipid extraction was repeated once by adding 450 µL of chloroform to the remaining cells in aqueous phase, and the lipid extracts were pooled into a single tube and dried in the SpeedVac under OH mode. Samples were stored at -80 ℃ until further analysis. Upper aqueous phase and cell pellet were dried in a SpeedVac under H_2_O mode. Total protein content was determined from the dried pellet using the Pierce® BCA Protein Assay Kit according to the manufacturer’s protocol.

Lipidomic analyses were conducted at LipidALL Technologies using a Shimadzu Nexera 20AD-HPLC/ExionLC-AD coupled with Sciex QTRAP 6500 PLUS. Separation of individual lipid classes of polar lipids by normal phase (NP)-HPLC was carried out using a TUP-HB silica column (i.d. 150×2.1 mm, 3 µm) with the following conditions: mobile phase A (chloroform: methanol:ammonium hydroxide, 89.5:10:0.5) and mobile phase B (chloroform:methanol:ammonium hydroxide:water, 55:39:0.5:5.5). MRM transitions were set up for comparative analysis of various polar lipids. Individual lipid species were quantified by referencing to spiked internal standards. d9-PC32:0(16:0/16:0), d9-PC36:1p(18:0p/18:1), d7-PE33:1(15:0/18:1), d9-PE36:1p(18:0p/18:1), d31-PS(d31-16:0/18:1), d7-PA33:1(15:0/18:1), d7-PG33:1(15:0/18:1), d7-PI33:1(15:0/18:1), C17-SL,d7-LPE18:1, C17-LPI, C17-LPA, C17-LPS were obtained from Avanti Polar Lipids. GM3-d18:1/18:0-d3 was purchased from Matreya LLC. Free fatty acids were quantitated using d31-16:0 (Sigma-Aldrich) and d8-20:4 (Cayman Chemicals).

### TAG and cholesterol quantification

Cells in six-well plates were lysed in 1% Triton X-100. Triacylglycerol was measured with a glycerol-phosphate-oxidase (GPO-PAP) colorimetric kit (BioSino Bio) and total cholesterol ester with a matched enzymatic colorimetric kit, read at 510 nm on a microplate reader. Total protein was determined in parallel by BCA assay (Thermo Fisher) and values are reported as mmol per g protein; the triacylglycerol measurements derive from n = 4 independent experiments.

### Image analysis and LD quantification

All quantification was performed in Fiji/ImageJ on raw, unprocessed images. For confocal LD counting, single mid-cell optical sections were background-subtracted (rolling ball, 50 px), thresholded with the Otsu algorithm applied identically to all conditions within an experiment, separated with a distance-transform watershed and measured with Analyze Particles (size filter 0.05–50 µm², circularity 0.3–1.0). Cell outlines were drawn manually from the DIC or ER-marker channel. Diameters were computed from the segmented area assuming a circular profile. LiveDrop puncta were segmented on the mCherry channel using the same pipeline with a 0.05–2 µm² size filter; puncta count in BLTP1-depleted cells include ring-shaped LiveDrop signal surrounding enlarged LDs.

### Colocalization analysis

For the two-color puncta analyses, channels were acquired sequentially and a punctum was scored as colocalized on the basis of a fixed overlap criterion applied identically to all conditions within an experiment.

### Structure prediction and interface confidence

Multi-chain predictions were run with AlphaFold 3 through the AlphaFold Server^73^. Two-chain BLTP1(1–67)–FIT2 predictions were additionally run with AlphaFold-Multimer in ColabFold using MMseqs2-generated MSAs, five models and three recycles, with monomer references from AlphaFold2. Interface confidence was assessed from ipTM and pTM. Predictions were treated as hypothesis-generating only and were not used as evidence of interaction.

### Statistics and reproducibility

Analyses were performed in GraphPad Prism 8.0.1. Data are mean ± SD unless stated otherwise. Two-group comparisons used two-tailed unpaired Student’s t-tests, and comparisons among three or more groups used ordinary one-way ANOVA with Tukey’s multiple-comparisons test; the test used for each panel is stated in the figure legend. Exact P values are reported in the figures and text; P < 0.05 was considered significant.

For lipidomics, class totals and individual species were compared by two-sided Welch’s *t*-tests on log2-transformed values, with effect sizes expressed as log2 ratios of geometric means with 95% confidence intervals, and multiplicity was controlled by the Benjamini–Hochberg procedure within each family (lipid classes, lipid species, the acyl-resolved PA transitions and the PI double-bond bins). Values below detection were replaced by half the minimum positive value of that feature before log transformation. Sample sizes were not predetermined by power analysis. No data were excluded. Randomization was not applicable; TEM tracing and adipocyte differentiation scoring were performed blinded to treatment condition, while other analyses were automated with condition-independent thresholds.

## Data availability

Reagents including plasmids and cell lines used in this study are available from the corresponding author on reasonable request.

## Acknowledgements

We thank School of Basic Medicine Innovation Research Center, Huazhong University of Science and Technology for assistance with imaging and flow cytometry. This work was supported by the National Natural Science Foundation of China (32541022, 32570822, 32371343, 32270779) and Shenzhen Medical Research Funds, General Program (B2602014).

## Author contributions

Q. Chu and W. Ji conceived the study and designed the experiments. Q. Chu, T. Zhou, Z. Zhou, and S. Zhang performed the imaging, biochemical and lipidomic experiments. Q. Chu, T. Zhou, Z. Zhou, S. Zhang and L. Deng performed and analyzed additional imaging, electron microscopy and adipogenic differentiation experiments. Q. Chu, Z. Zhou and W. Ji analyzed and interpreted the data. W. Ji wrote the manuscript with input from all authors. All authors read and approved the final manuscript.

**Correspondence and requests for materials** should be addressed to W.-K.J.

---

**Extended Data Fig. 1.**
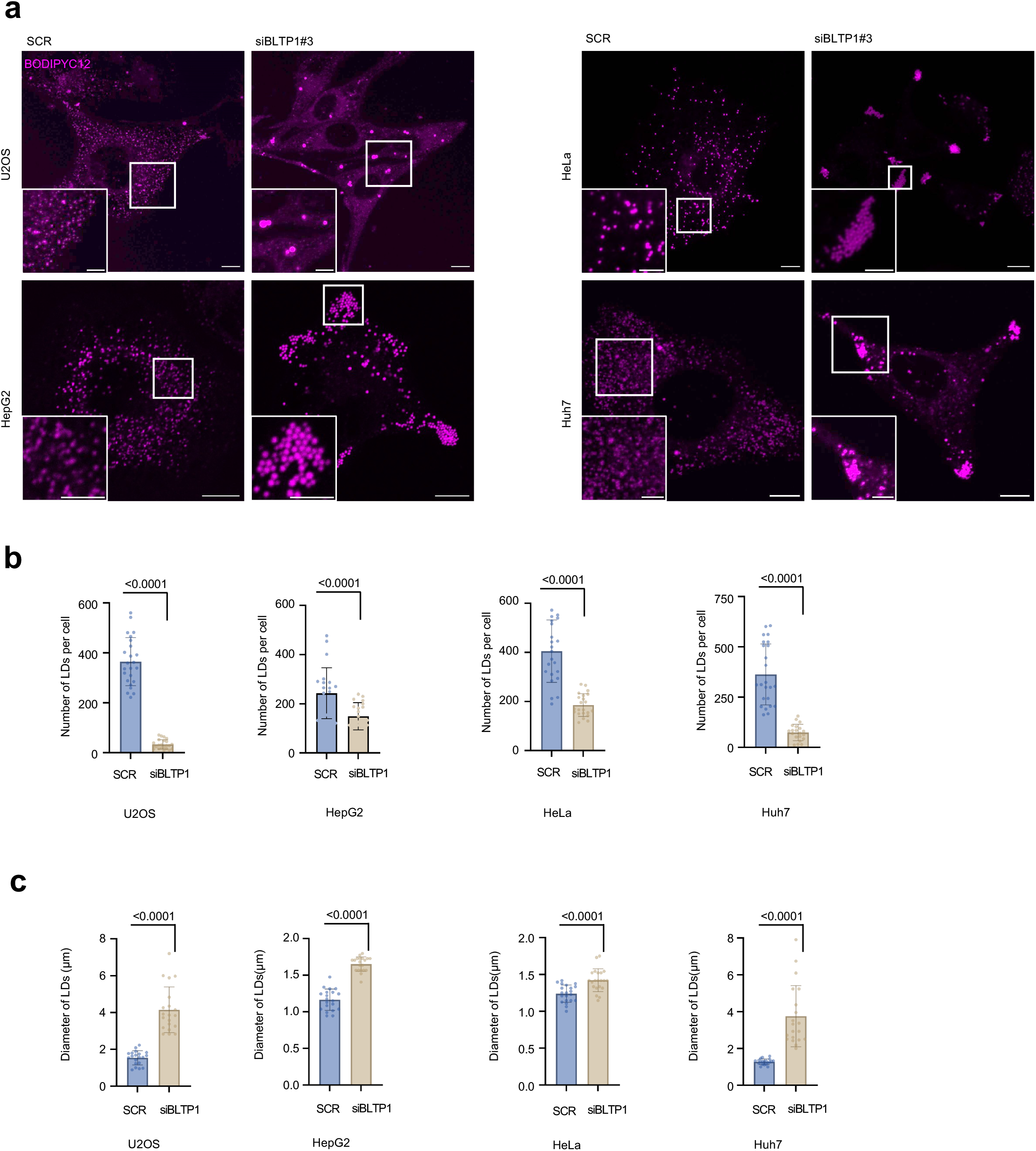
The BLTP1 LD phenotype is conserved across cell lines. **a**, Representative BODIPY images of OA-loaded U2OS, HepG2, HeLa and Huh7 cells treated with SCR or siBLTP1 with insets. **b**, quantification of LD number from cells as in (**a**) from three independent experiments. Mean ± SD. Two-tailed unpaired Student’s t-tests. **c**, LD diameter from cells as in (**a**) from three independent experiments. Mean ± SD. Two-tailed unpaired Student’s t-tests. Scale bars: 10 μm in whole-cell image and 2 μm in insets.

**Extended Data Fig. 2.**
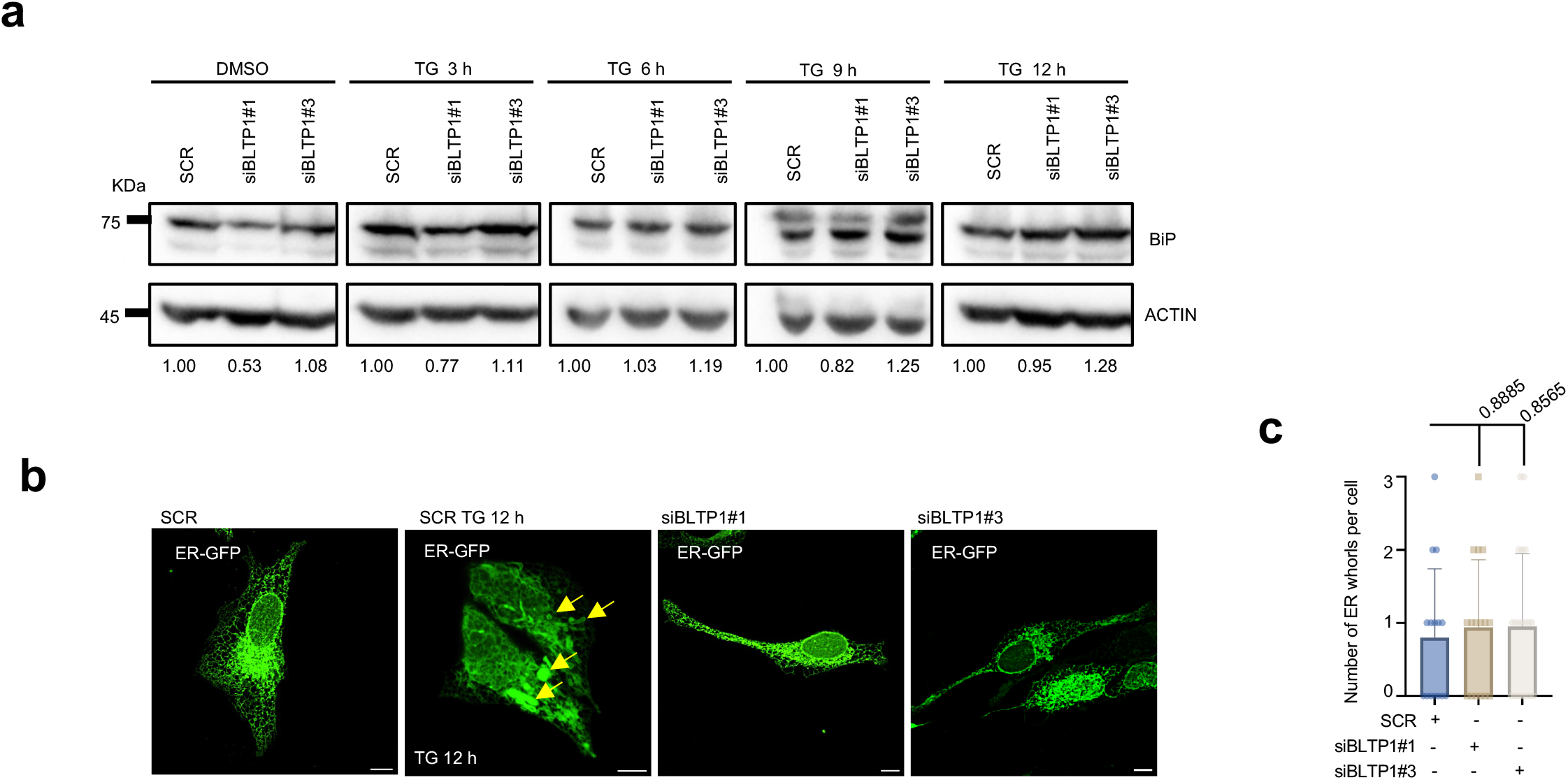
BLTP1 depletion does not induce ER stress. **a,** Representative BiP immunoblots of U2OS cells transfected with SCR, siBLTP1#1 or siBLTP1#3 and treated with DMSO or thapsigargin (TG; 2 μM for 3, 6, 9 or 12 h); actin, loading control. Numbers below the lanes are BiP signal normalized to actin. **b,** Representative confocal images of live U2OS cells expressing ER-GFP (ER luminal marker) treated with SCR or BLTP1 siRNA; SCR cells treated with thapsigargin (2 μM, 12 h) serve as the positive control. Yellow arrows denote potential ER whorls. **c,** Quantification of ER whorls per cell section scored from confocal microscopy in (**b**) from three independent experiments. Mean ± SD. Ordinary one-way ANOVA with Tukey’s multiple-comparisons test. Scale bars: 10 μm in whole-cell image in (c).

**Extended Data Fig. 3.**
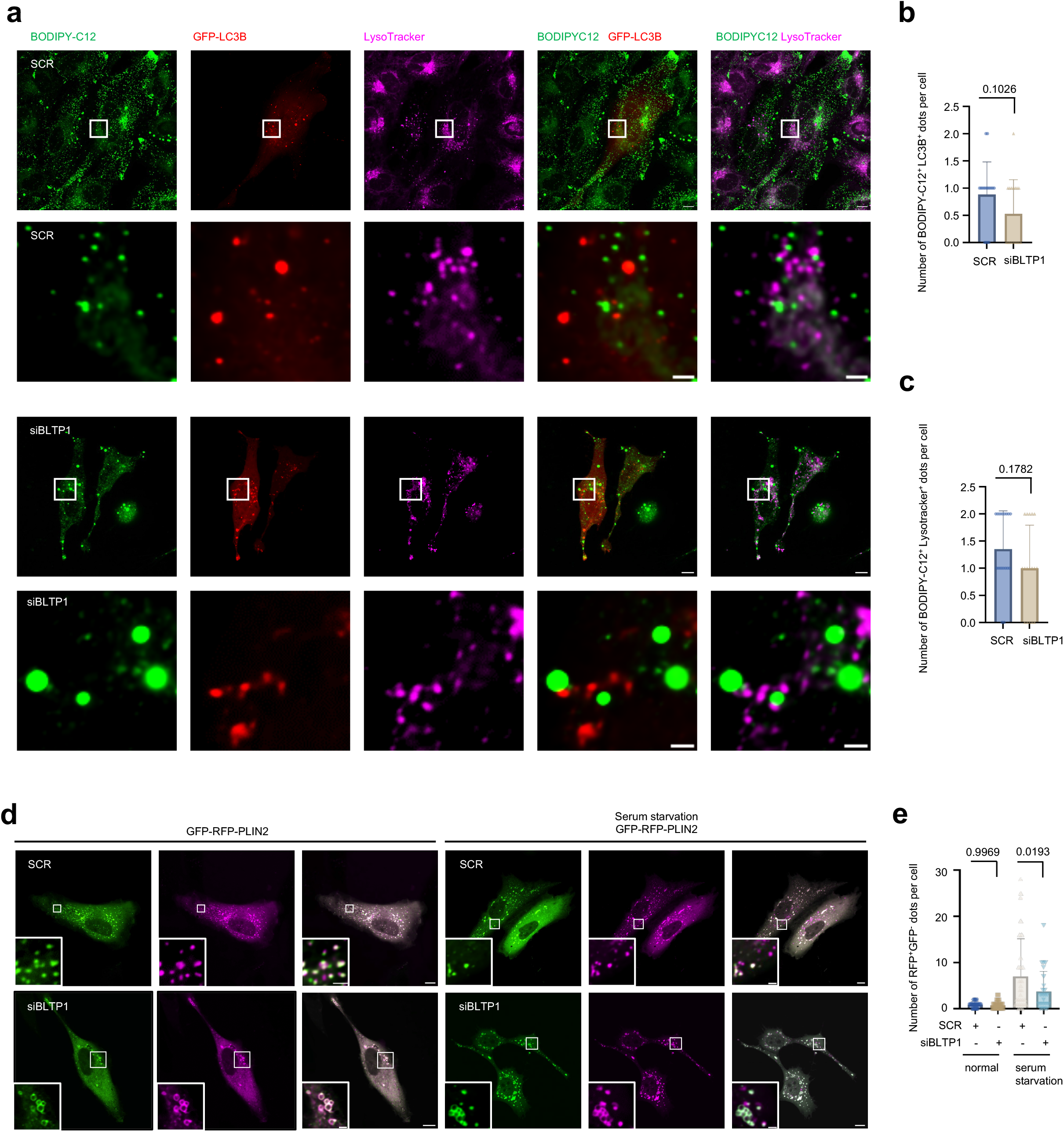
LD–autophagosome and LD–lysosome contacts and LD turnover are largely unaffected by BLTP1 depletion. **a,** Representative live-cell confocal images from BODIPY-C12 (green) and LysoTracker (magenta)-stained U2OS cells expressing GFP-LC3B (red) treated with SCR and BLTP1 siRNAs. **b,** Quantification of number of BODIPY-C12^+^ LC3B^+^ dots per cell as in (**a**) from three independent experiments. Mean ± SD. Two-tailed unpaired Student’s t-tests. **c,** Quantification of number of BODIPY-C12^+^ LysoTracker^+^ dots per cell as in (**a**) from three independent experiments. Mean ± SD. Two-tailed unpaired Student’s t-tests. **d,** Representative live-cell confocal images of U2OS cells expressing GFP-RFP-PLIN2 under normal (left panel) or serum starvation (6 h; right panel) with insets on the bottom. **e,** Quantification of number of RFP^+^GFP^−^dots per cell as in (**d**) from 3 independent experiments with at least 15 cells quantified per group. Two-tailed unpaired Student’s t test. Mean ± SD. Scale bar, 10 μm in the whole cell images and 2 μm in the insets.

**Extended Data Fig. 4.**
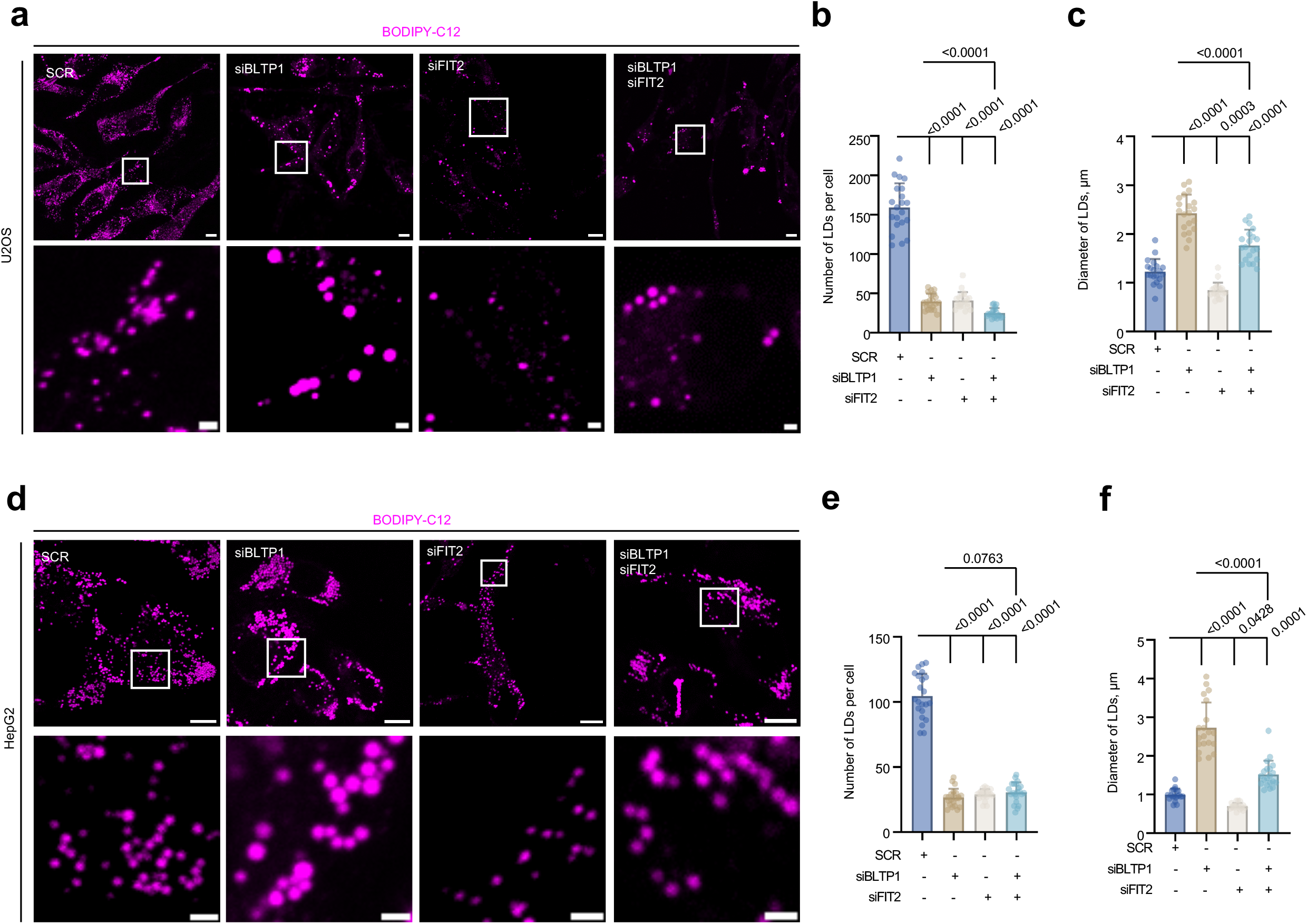
Co-depletion of FIT2 suppresses the LD enlargement caused by BLTP1 depletion. **a**, Representative live-cell confocal BODIPY images of OA-loaded U2OS cells treated with SCR, siFIT2, siBLTP1 or siBLTP1 + siFIT2 with insets on the bottom. **b**, LD number per cell in (**a**) for SCR (21 cells), siBLTP1 (21 cells), siFIT2 (21 cells) or double knockdown (21 cells) from three independent experiments. Mean ± SD. Ordinary one-way ANOVA with Tukey’s multiple-comparisons test. **c**, LD diameter in cells from (**a**) from three independent experiments. Mean ± SD. Ordinary one-way ANOVA with Tukey’s multiple-comparisons test. **d**, Representative live-cell confocal BODIPY images of OA-loaded HepG2 cells treated with SCR, siFIT2, siBLTP1 or siBLTP1 + siFIT2 with insets on the bottom. **e**, LD number per cell in (**d**) for SCR (22 cells), siBLTP1 (22 cells), siFIT2 (22 cells) or double knockdown (22 cells) from three independent experiments. Mean ± SD. Ordinary one-way ANOVA with Tukey’s multiple-comparisons test. **f**, LD diameter in cells from (**d**) from three independent experiments. Mean ± SD. Ordinary one-way ANOVA with Tukey’s multiple-comparisons test. Scale bar, 10 μm in the whole cell image and 2 μm in the insets.

**Extended Data Fig. 5.**
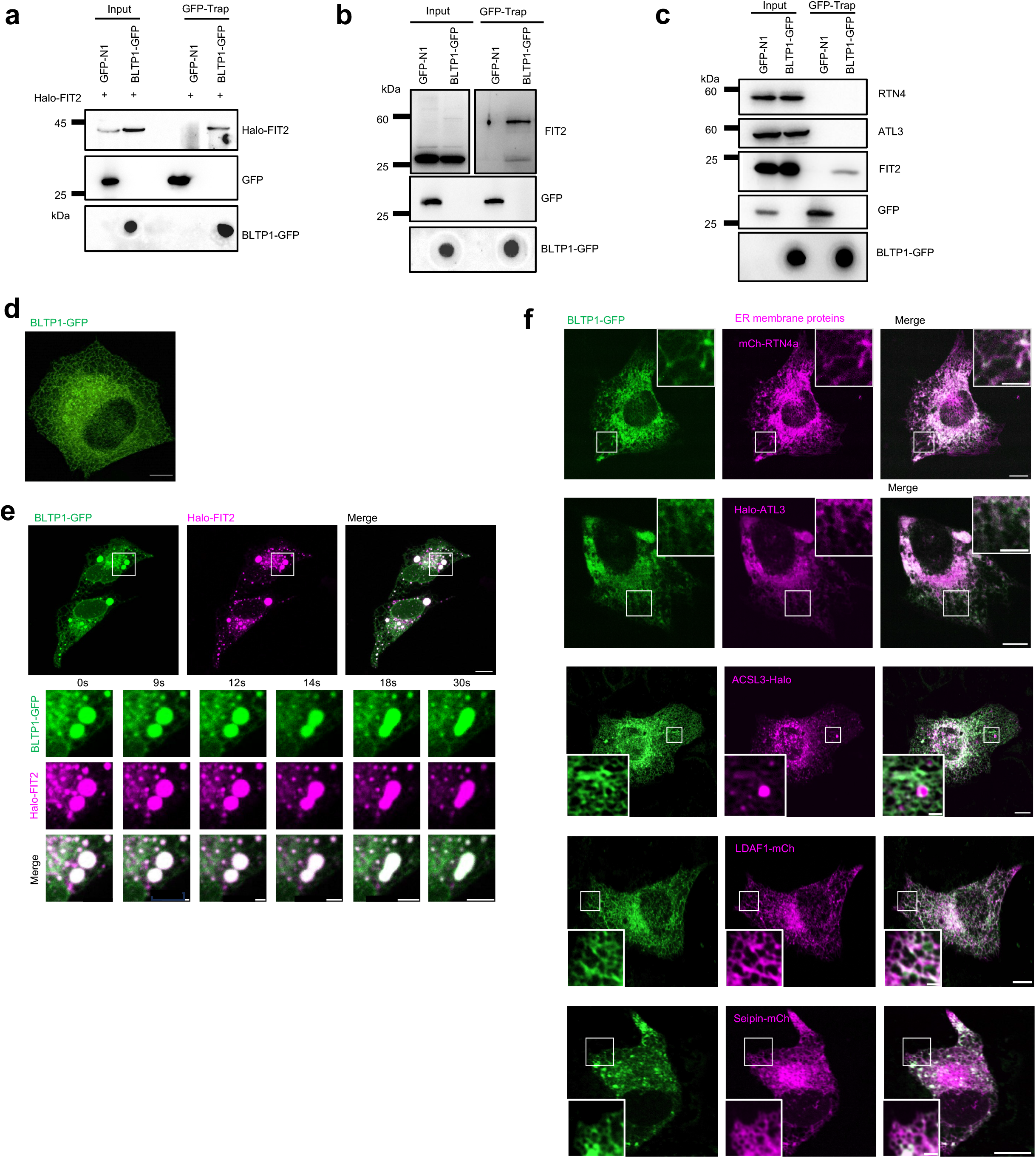
BLTP1 colocalizes with FIT2 but not with other ER or LD-biogenesis proteins. **a,** Representative GFP-Trap assays of HEK293 cells expressing Halo-FIT2 along with GFP empty vector or BLTP1-GFP. Dot blots were used to detect BLTP1-GFP. **b,** Representative GFP-Trap assays of HEK293 cells expressing GFP empty vector or BLTP1-GFP. Endogenous FIT2 was detected in this assay. Dot blots were used to detect BLTP1-GFP. **c,** Representative GFP-Trap assays of HEK293 cells expressing GFP empty vector or BLTP1-GFP. ER membrane proteins such as RTN4, ATL3 and FIT2 were detected at endogenous levels in this assay. Dot blots were used to detect BLTP1-GFP. **d**, Representative live-cell confocal image of U2OS cells expressing BLTP1-GFP alone. **e**, Representative live-cell confocal image of U2OS cells expressing BLTP1-GFP (green) and Halo-FIT2 (magenta) with insets showing dynamics of BLTP1–GFP and Halo–FIT2 puncta. Time in second. **f**, Representative live-cell confocal image of U2OS cells expressing BLTP1-GFP (green) and ER membrane proteins (magenta; mCh-RTN4, Halo-ATL3, ACSL3-Halo, LDAF1-mCh and Seipin-mCh) with insets. Scale bar, 10 μm in the whole cell image and 2 μm in the insets.

**Extended Data Fig. 6.**
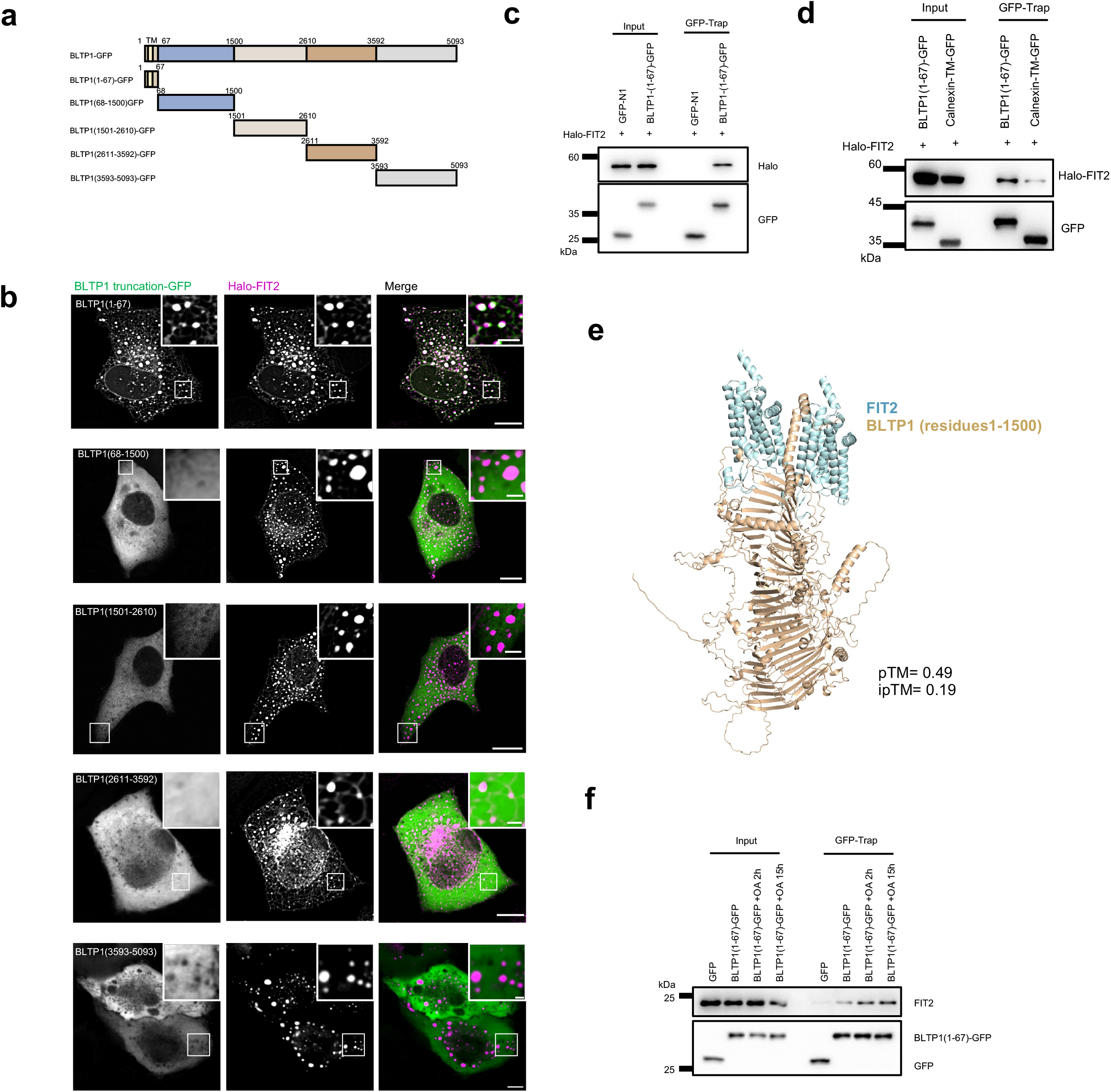
The BLTP1 N-terminal region is necessary and sufficient for FIT2 binding. **a**, The schematic diagram of BLTP1 and truncations. **b**, Representative images of live U2OS cells expressing Halo-FIT2 (magenta) and BLTP1 truncations (green) with insets. **c**, Representative GFP-Trap assays demonstrate an interaction between Halo-FIT2 and BLTP1(1-67)-GFP. **d**, Representative GFP-Trap assays of HEK293 cells expressing Halo-FIT2 along with BLTP1(1-67)-GFP or Calnexin-TM-GFP. **e**, AlphaFold-Multimer model of FIT2 (cyan) with BLTP1(1–1500) (tan); no confident interface was predicted (pTM = 0.49; ipTM = 0.19). **f**, Representative GFP-Trap assays of endogenous FIT2 with BLTP1(1-67)–GFP in the presence or absence of OA treatment. Scale bar, 10 μm in the whole cell image and 2 μm in the insets

**Extended Data Fig. 7.**
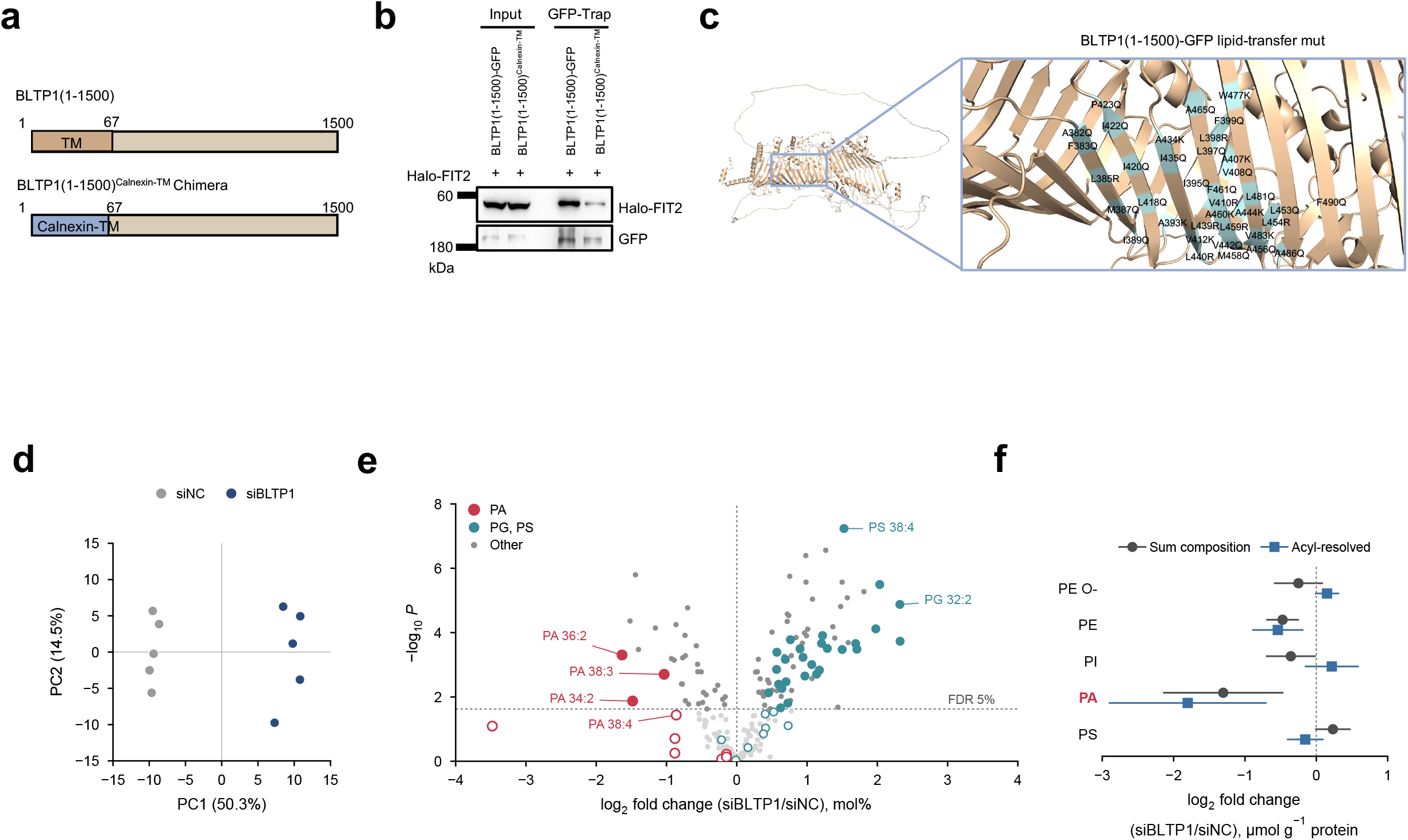
The BLTP1 mutants and remodeling of the LD-fraction lipidome in BLTP1-depleted U2OS cells. **a**, Domain schematics of BLTP1(1-1500)^Calnexin-TM^ Chimera, in which residues 1–67 of BLTP1 are replaced by the transmembrane segment of calnexin (Calnexin-TM). **b**, Representative GFP-Trap assays comparing Halo-FIT2 binding to BLTP1(1-1500)-GFP and to the BLTP1(1-1500)^Calnexin-TM^ chimera, immunoblotted for Halo and GFP. **c**, The AlphaFold3-predicted structures of BLTP1(1-1500)-GFP mut carrying 37 groove substitutions (A382Q, F383Q, L385R, M387Q, I389Q, A393K, I395Q, L397Q, L398R, F399Q, A407K, V408Q, V410R, V412K, L418Q, I420Q, I422Q, P423Q, A434K, I435Q, L439R, L440R, V442Q, A444K, L453Q, L454R, A456Q, M458Q, L459R, A460K, F461Q, A465Q, W477K, L481Q, V483K, A486Q, F490Q), designed after the RBG groove definitions and ATG2 cavity-mutagenesis strategy^51,57,60^. **d**, Principal component analysis of autoscaled log2 mol% values of 199 polar-lipid species in LD fractions. **e**, Volcano plot of the 199 species (mol%). Filled symbols, *q* < 0.05; the dashed line marks the 5% false-discovery threshold. Red, PA; teal, PG and PS; grey, other classes. **f**, Class sums per unit protein, calculated separately from the sum-composition and the acyl-resolved transitions of the targeted method. Points show log2 ratios of geometric means and lines 95% confidence intervals. The two channels were not pooled. For **d**–**f**, n = 5 LD preparations per group, each from an independent transfection, and *P* values are from two-sided Welch’s *t*-tests on log2-transformed values. Samples are as in Fig. 5. Source data are provided as Source Data Extended Data Fig. 5.

**Extended Data Fig. 8.**
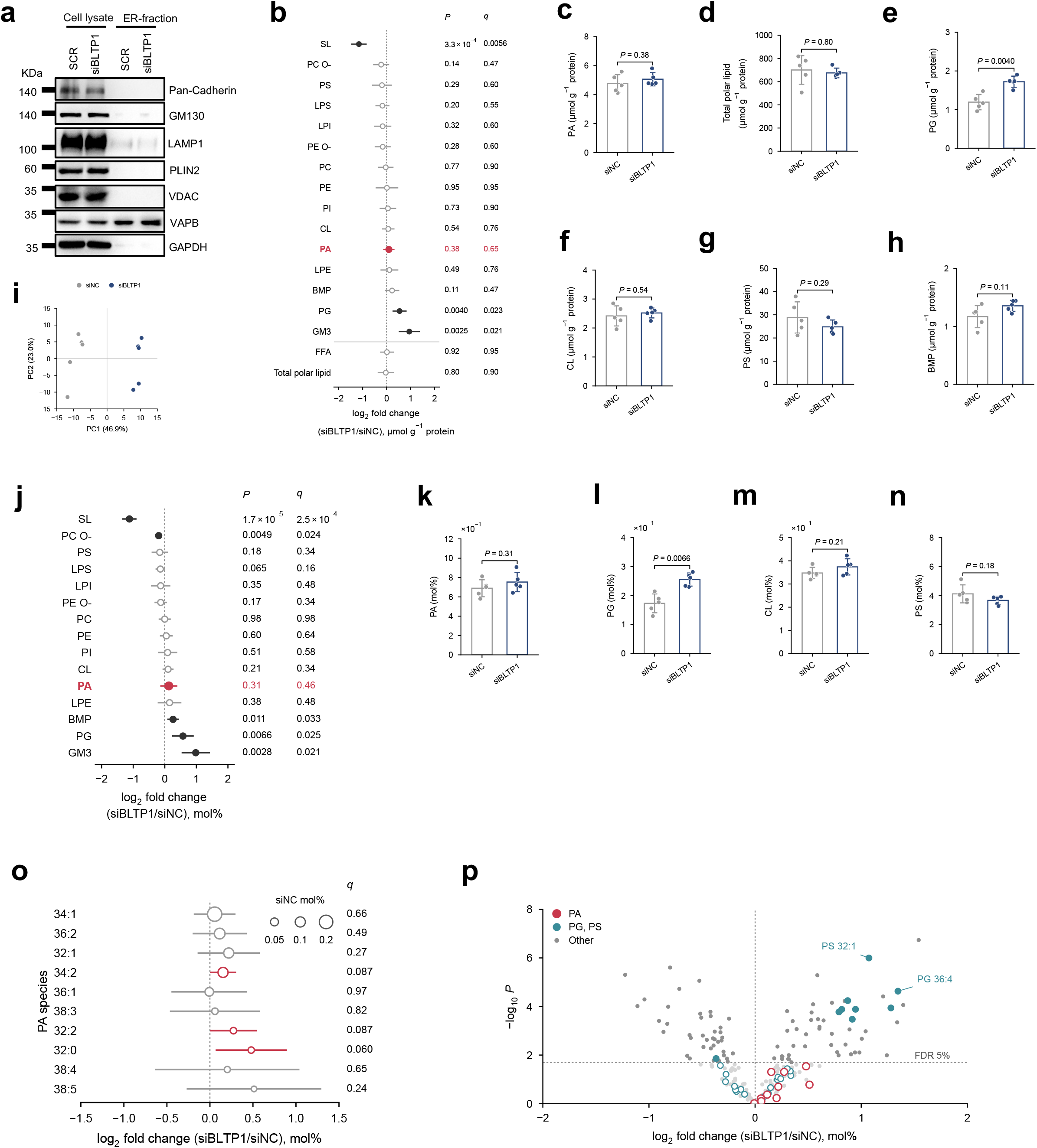
BLTP1 depletion does not change PA in the ER. **a**, Representative immunoblots of total cell lysates and ER fractions from SCR and siBLTP1 U2OS cells, loaded with OA for 16 h. The markers are Pan-cadherin (plasma membrane), GM130 (Golgi), LAMP1 (lysosome), PLIN2 (LD), VDAC (mitochondria), VAPB (ER) and GAPDH (cytosol). **b**, Class-level change in ER-fraction lipids normalized to fraction protein, displayed as in Fig. 5b (BH across the 17 features shown). **c**–**h**, PA (**c**), total polar lipid (**d**), PG (**e**), CL (**f**), PS (**g**) and BMP (**h**) per unit fraction protein. **i**, Principal component analysis of autoscaled log2 mol% values of 208 polar-lipid species. **j**, As in (**b)**, with classes expressed as mol% of total measured polar lipid (BH across 15 classes). **k**–**n**, PA (**k**), PG (**l**), CL (**m**) and PS (**n**) as mol% of polar lipid. **o**, Change in PA species expressed as mol%, displayed as in Fig. 5k (BH across 208 species). **p**, Volcano plot of the 208 species (mol%), displayed as in Extended Data Fig. 7e. Mean ± SD, and dots individual ER preparations. n = 5 per group. *P* values are from two-sided Welch’s *t*-tests on log2-transformed values. LPS, lyso-PS; other abbreviations as in Fig. 5.

**Extended Data Fig. 9.**
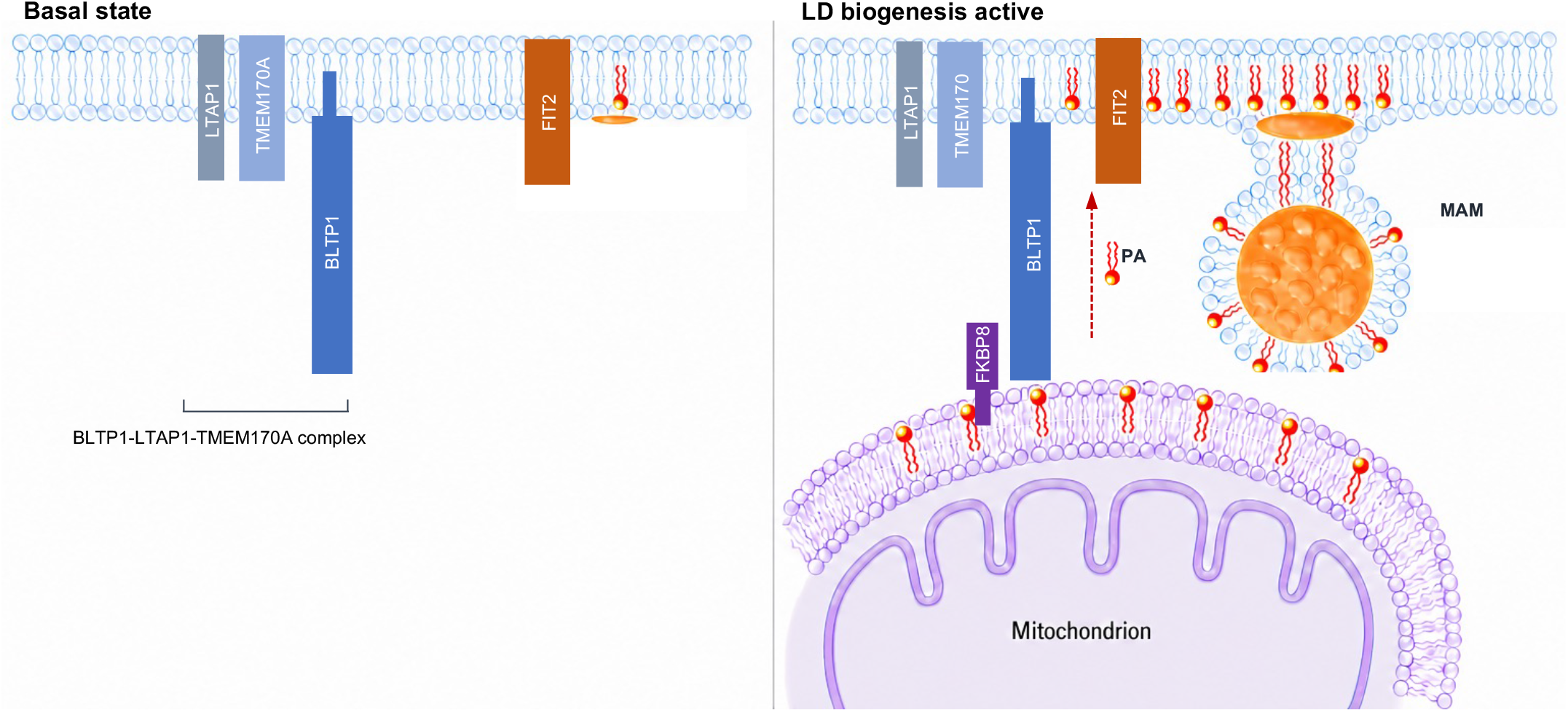
Proposed model of BLTP1-dependent PA supply at LD formation sites. Left, basal state: BLTP1 resides in the ER in a complex with LTAP1 and TMEM170A, separate from FIT2. Right, active LD biogenesis: BLTP1 engages FIT2 through its N-terminal transmembrane segment, and the BLTP1–FIT2 module maintains local PA availability at the LD formation site, which determines how efficiently nascent LDs form and how large they grow. The C-terminal region of BLTP1 binds the outer mitochondrial membrane protein FKBP8, as characterized in a companion study^67^, placing the bridge at mitochondria-associated ER membranes (MAM). The dashed arrow denotes a hypothetical route of PA transfer from the outer mitochondrial membrane to the ER; the source and direction of PA flux were not tested in this study.

## References

1 Walther, T. C., Chung, J. & Farese, R. V. Lipid Droplet Biogenesis. Annual Review of Cell and Developmental Biology 33, 491–510, doi:10.1146/annurev-cellbio-100616-060608 (2017).

2 Olzmann, J. A. & Carvalho, P. Dynamics and functions of lipid droplets. Nature Reviews Molecular Cell Biology 20, 137–155, doi:10.1038/s41580-018-0085-z (2018).

3 Thiam, A. R., Farese Jr, R. V. & Walther, T. C. The biophysics and cell biology of lipid droplets. Nature Reviews Molecular Cell Biology 14, 775–786, doi:10.1038/nrm3699 (2013).

4 Ben M’barek, K., et al. ER Membrane Phospholipids and Surface Tension Control Cellular Lipid Droplet Formation. Developmental Cell 41, 591–604.e597, doi:10.1016/j.devcel.2017.05.012 (2017).

5 Choudhary, V. et al. Architecture of Lipid Droplets in Endoplasmic Reticulum Is Determined by Phospholipid Intrinsic Curvature. Current Biology 28, 915–926.e919, doi:10.1016/j.cub.2018.02.020 (2018).

6 Chorlay, A. et al. Membrane Asymmetry Imposes Directionality on Lipid Droplet Emergence from the ER. Developmental Cell 50, 25–42.e27, doi:10.1016/j.devcel.2019.05.003 (2019).

7 Santinho, A. et al. Membrane Curvature Catalyzes Lipid Droplet Assembly. Current Biology 30, 2481–2494.e2486, doi:10.1016/j.cub.2020.04.066 (2020).

8 Wilfling, F. et al. Triacylglycerol Synthesis Enzymes Mediate Lipid Droplet Growth by Relocalizing from the ER to Lipid Droplets. Developmental Cell 24, 384–399, doi:10.1016/j.devcel.2013.01.013 (2013).

9 Kassan, A. et al. Acyl-CoA synthetase 3 promotes lipid droplet biogenesis in ER microdomains. Journal of Cell Biology 203, 985–1001, doi:10.1083/jcb.201305142 (2013).

10 Wang, H. et al. Seipin is required for converting nascent to mature lipid droplets. eLife 5, doi:10.7554/elife.16582 (2016).

11 Magré, J. et al. Identification of the gene altered in Berardinelli–Seip congenital lipodystrophy on chromosome 11q13. Nature Genetics 28, 365–370, doi:10.1038/ng585 (2001).

12 Szymanski, K. M. et al. The lipodystrophy protein seipin is found at endoplasmic reticulum lipid droplet junctions and is important for droplet morphology. Proceedings of the National Academy of Sciences 104, 20890–20895, doi:10.1073/pnas.0704154104 (2007).

13 Sui, X. et al. Cryo–electron microscopy structure of the lipid droplet–formation protein seipin. Journal of Cell Biology 217, 4080–4091, doi:10.1083/jcb.201809067 (2018).

14 Klug, Y. A. et al. Mechanism of lipid droplet formation by the yeast Sei1/Ldb16 Seipin complex. Nature Communications 12, doi:10.1038/s41467-021-26162-6 (2021).

15 Arlt, H. et al. Seipin forms a flexible cage at lipid droplet formation sites. Nature Structural & Molecular Biology 29, 194–202, doi:10.1038/s41594-021-00718-y (2022).

16 Yan, R. et al. Human SEIPIN Binds Anionic Phospholipids. Developmental Cell 47, 248–256.e244, doi:10.1016/j.devcel.2018.09.010 (2018).

17 Prasanna, X. et al. Seipin traps triacylglycerols to facilitate their nanoscale clustering in the endoplasmic reticulum membrane. PLOS Biology 19, e3000998, doi:10.1371/journal.pbio.3000998 (2021).

18 Castro, I. et al. Promethin Is a Conserved Seipin Partner Protein. Cells 8, 268, doi:10.3390/cells8030268 (2019).

19 Chung, J. et al. LDAF1 and Seipin Form a Lipid Droplet Assembly Complex. Developmental Cell 51, 551–563.e557, doi:10.1016/j.devcel.2019.10.006 (2019).

20 Salo, V. T. et al. Seipin regulates ER–lipid droplet contacts and cargo delivery. The EMBO Journal 35, 2699–2716, doi:10.15252/embj.201695170 (2016).

21 Salo, V. T. et al. Seipin Facilitates Triglyceride Flow to Lipid Droplet and Counteracts Droplet Ripening via Endoplasmic Reticulum Contact. Developmental Cell 50, 478–493.e479, doi:10.1016/j.devcel.2019.05.016 (2019).

22 Li, C. et al. Adipogenin promotes the development of lipid droplets by binding a dodecameric seipin complex. Science 390, doi:10.1126/science.adr9755 (2025).

23 Krahmer, N. et al. Phosphatidylcholine Synthesis for Lipid Droplet Expansion Is Mediated by Localized Activation of CTP:Phosphocholine Cytidylyltransferase. Cell Metabolism 14, 504–515, doi:10.1016/j.cmet.2011.07.013 (2011).

24 Han, G.-S., Wu, W.-I. & Carman, G. M. The Saccharomyces cerevisiae Lipin Homolog Is a Mg2+-dependent Phosphatidate Phosphatase Enzyme*. Journal of Biological Chemistry 281, 9210–9218, doi:10.1074/jbc.m600425200 (2006).

25 Cases, S. et al. Identification of a gene encoding an acyl CoA:diacylglycerol acyltransferase, a key enzyme in triacylglycerol synthesis. Proceedings of the National Academy of Sciences 95, 13018–13023, doi:10.1073/pnas.95.22.13018 (1998).

26 Fei, W. et al. A Role for Phosphatidic Acid in the Formation of “Supersized” Lipid Droplets. PLoS Genetics 7, e1002201, doi:10.1371/journal.pgen.1002201 (2011).

27 Prinz, W. A., Toulmay, A. & Balla, T. The functional universe of membrane contact sites. Nature Reviews Molecular Cell Biology 21, 7–24, doi:10.1038/s41580-019-0180-9 (2019).

28 Chung, J. et al. PI4P/phosphatidylserine countertransport at ORP5- and ORP8-mediated ER–plasma membrane contacts. Science 349, 428–432, doi:10.1126/science.aab1370 (2015).

29 Du, X. et al. ORP5 localizes to ER–lipid droplet contacts and regulates the level of PI(4)P on lipid droplets. Journal of Cell Biology 219, doi:10.1083/jcb.201905162 (2019).

30 Guyard, V. et al. ORP5 and ORP8 orchestrate lipid droplet biogenesis and maintenance at ER–mitochondria contact sites. Journal of Cell Biology 221, doi:10.1083/jcb.202112107 (2022).

31 Zhou, T. et al. Sec14L6 is a phosphoinositide transporter that regulates phosphoinositide homeostasis and biogenesis of lipid droplets. Nature Communications 16, doi:10.1038/s41467-025-65540-2 (2025).

32 Freyre, C. A. C., Rauher, P. C., Ejsing, C. S. & Klemm, R. W. MIGA2 Links Mitochondria, the ER, and Lipid Droplets and Promotes De Novo Lipogenesis in Adipocytes. Molecular Cell 76, 811–825.e814, doi:10.1016/j.molcel.2019.09.011 (2019).

33 Yeshaw, W. M. et al. Human VPS13A is associated with multiple organelles and influences mitochondrial morphology and lipid droplet motility. eLife 8, doi:10.7554/elife.43561 (2019).

34 Kadereit, B. et al. Evolutionarily conserved gene family important for fat storage. Proceedings of the National Academy of Sciences 105, 94–99, doi:10.1073/pnas.0708579105 (2008).

35 Gross, D. A., Snapp, E. L. & Silver, D. L. Structural Insights into Triglyceride Storage Mediated by Fat Storage-Inducing Transmembrane (FIT) Protein 2. PLoS ONE 5, e10796, doi:10.1371/journal.pone.0010796 (2010).

36 Gross, D. A., Zhan, C. & Silver, D. L. Direct binding of triglyceride to fat storage-inducing transmembrane proteins 1 and 2 is important for lipid droplet formation. Proceedings of the National Academy of Sciences 108, 19581–19586, doi:10.1073/pnas.1110817108 (2011).

37 Choudhary, V., Ojha, N., Golden, A. & Prinz, W. A. A conserved family of proteins facilitates nascent lipid droplet budding from the ER. Journal of Cell Biology 211, 261–271, doi:10.1083/jcb.201505067 (2015).

38 Hayes, M. J. et al. Fat storage-inducing transmembrane (FIT or FITM) proteins are related to lipid phosphatase/phosphotransferase enzymes. Microbial Cell 5, 88–103, doi:10.15698/mic2018.02.614 (2018).

39 Moir, R. D., Gross, D. A., Silver, D. L. & Willis, I. M. SCS3 and YFT2 Link Transcription of Phospholipid Biosynthetic Genes to ER Stress and the UPR. PLoS Genetics 8, e1002890, doi:10.1371/journal.pgen.1002890 (2012).

40 Becuwe, M. et al. FIT2 is an acyl–coenzyme A diphosphatase crucial for endoplasmic reticulum homeostasis. Journal of Cell Biology 219, doi:10.1083/jcb.202006111 (2020).

41 Chen, F. et al. FIT2 organizes lipid droplet biogenesis with ER tubule-forming proteins and septins. Journal of Cell Biology 220, doi:10.1083/jcb.201907183 (2021).

42 Graff, J. & Schneiter, R. FIT2 proteins and lipid droplet emergence, an interplay between phospholipid synthesis, surface tension, and membrane curvature. Frontiers in Cell and Developmental Biology 12, 1422032, doi:10.3389/fcell.2024.1422032 (2024).

43 Wang, G. et al. Fat storage-inducing transmembrane proteins: beyond mediating lipid droplet formation. Cellular & Molecular Biology Letters 27, doi:10.1186/s11658-022-00391-z (2022).

44 Miranda, D. A. et al. Fat Storage-inducing Transmembrane Protein 2 Is Required for Normal Fat Storage in Adipose Tissue. Journal of Biological Chemistry 289, 9560–9572, doi:10.1074/jbc.m114.547687 (2014).

45 Goh, V. J. et al. Postnatal Deletion of Fat Storage-inducing Transmembrane Protein 2 (FIT2/FITM2) Causes Lethal Enteropathy. Journal of Biological Chemistry 290, 25686–25699, doi:10.1074/jbc.m115.676700 (2015).

46 Zheng, X. et al. Destabilization of β Cell FIT2 by saturated fatty acids alter lipid droplet numbers and contribute to ER stress and diabetes. Proceedings of the National Academy of Sciences 119, doi:10.1073/pnas.2113074119 (2022).

47 Bond, L. M. et al. Fitm2 is required for ER homeostasis and normal function of murine liver. Journal of Biological Chemistry 299, 103022, doi:10.1016/j.jbc.2023.103022 (2023).

48 Seco, C. Z. et al. A homozygousFITM2mutation causes a deafness-dystonia syndrome with motor regression and signs of ichthyosis and sensory neuropathy. Disease Models & Mechanisms, doi:10.1242/dmm.026476 (2016).

49 Riedhammer, K. M., Leszinski, G. S., Andres, S., Strobl-Wildemann, G. & Wagner, M. First replication that biallelic variants in FITM2 cause a complex deafness-dystonia syndrome. Movement Disorders 33, 1665–1666, doi:10.1002/mds.27481 (2018).

50 Braschi, B., Bruford, E. A., Cavanagh, A. T., Neuman, S. D. & Bashirullah, A. The bridge-like lipid transfer protein (BLTP) gene group: introducing new nomenclature based on structural homology indicating shared function. Human Genomics 16, doi:10.1186/s40246-022-00439-3 (2022).

51 Levine, T. P. Sequence Analysis and Structural Predictions of Lipid Transfer Bridges in the Repeating Beta Groove (RBG) Superfamily Reveal Past and Present Domain Variations Affecting Form, Function and Interactions of VPS13, ATG2, SHIP164, Hobbit and Tweek. Contact 5, 251525642211343, doi:10.1177/25152564221134328 (2022).

52 Neuman, S. D., Levine, T. P. & Bashirullah, A. A novel superfamily of bridge-like lipid transfer proteins. Trends in Cell Biology 32, 962–974, doi:10.1016/j.tcb.2022.03.011 (2022).

53 Hanna, M., Guillén-Samander, A. & De Camilli, P. RBG Motif Bridge-Like Lipid Transport Proteins: Structure, Functions, and Open Questions. Annual Review of Cell and Developmental Biology 39, 409–434, doi:10.1146/annurev-cellbio-120420-014634 (2023).

54 Kumar, N. et al. VPS13A and VPS13C are lipid transport proteins differentially localized at ER contact sites. Journal of Cell Biology 217, 3625–3639, doi:10.1083/jcb.201807019 (2018).

55 Li, P., Lees, J. A., Lusk, C. P. & Reinisch, K. M. Cryo-EM reconstruction of a VPS13 fragment reveals a long groove to channel lipids between membranes. Journal of Cell Biology 219, doi:10.1083/jcb.202001161 (2020).

56 Adlakha, J., Hong, Z., Li, P. & Reinisch, K. M. Structural and biochemical insights into lipid transport by VPS13 proteins. Journal of Cell Biology 221, e202202030, doi:10.1083/jcb.202202030 (2022).

57 Osawa, T. et al. Atg2 mediates direct lipid transfer between membranes for autophagosome formation. Nature Structural & Molecular Biology 26, 281–288, doi:10.1038/s41594-019-0203-4 (2019).

58 Valverde, D. P. et al. ATG2 transports lipids to promote autophagosome biogenesis. Journal of Cell Biology 218, 1787–1798, doi:10.1083/jcb.201811139 (2019).

59 Maeda, S., Otomo, C. & Otomo, T. The autophagic membrane tether ATG2A transfers lipids between membranes. eLife 8, doi:10.7554/elife.45777 (2019).

60 Kang, Y. et al. Structural basis of lipid transfer by a bridge-like lipid-transfer protein. Nature 642, 242–249, doi:10.1038/s41586-025-08918-y (2025).

61 Wang, C. et al. A conserved megaprotein-based molecular bridge critical for lipid trafficking and cold resilience. Nature Communications 13, doi:10.1038/s41467-022-34450-y (2022).

62 Toulmay, A. et al. Vps13-like proteins provide phosphatidylethanolamine for GPI anchor synthesis in the ER. Journal of Cell Biology 221, doi:10.1083/jcb.202111095 (2022).

63 Verstreken, P. et al. Tweek, an Evolutionarily Conserved Protein, Is Required for Synaptic Vesicle Recycling. Neuron 63, 203–215, doi:10.1016/j.neuron.2009.06.017 (2009).

64 Rocha-Roa, C. et al. TMEM170 proteins are lipid scramblases associated with bridge-type lipid transporters BLTP1/Csf1. Nature Structural & Molecular Biology 33, 215–219, doi:10.1038/s41594-025-01716-0 (2025).

65 Gueneau, L. et al. KIAA1109 Variants Are Associated with a Severe Disorder of Brain Development and Arthrogryposis. The American Journal of Human Genetics 102, 116–132, doi:10.1016/j.ajhg.2017.12.002 (2018).

66 Liu, Y., Ye, Q., Ma, D. K., Rothermel, B. A. & Lin, W. Bltp1 Is Required for Survival and Normal Development and Function of the Neuromuscular Junction in Mice. The Journal of Neuroscience 45, e0029252025, doi:10.1523/jneurosci.0029-25.2025 (2025).

67 Chu, Q. & Ji, W. BLTP1-dependent phospholipid efflux prevents mitochondrial lipid overload and apoptosis at FKBP8-tethered ER-mitochondrial contact. Preprint at bioRxiv 10.1101/2025.09.30.679455 (2025).

68 Rambold, A. S., Cohen, S. & Lippincott-Schwartz, J. Fatty Acid Trafficking in Starved Cells: Regulation by Lipid Droplet Lipolysis, Autophagy, and Mitochondrial Fusion Dynamics. Developmental Cell 32, 678–692, doi:10.1016/j.devcel.2015.01.029 (2015).

69 Brasaemle, D. L. et al. Adipose differentiation-related protein is an ubiquitously expressed lipid storage droplet-associated protein. Journal of Lipid Research 38, 2249–2263, doi:10.1016/s0022-2275(20)34939-7 (1997).

70 Los, G. V. et al. HaloTag: A Novel Protein Labeling Technology for Cell Imaging and Protein Analysis. ACS Chemical Biology 3, 373–382, doi:10.1021/cb800025k (2008).

71 Grimm, J. B. et al. A general method to improve fluorophores for live-cell and single-molecule microscopy. Nature Methods 12, 244–250, doi:10.1038/nmeth.3256 (2015).

72 Sun, Y., Day, R. N. & Periasamy, A. Investigating protein-protein interactions in living cells using fluorescence lifetime imaging microscopy. Nature Protocols 6, 1324–1340, doi:10.1038/nprot.2011.364 (2011).

73 Abramson, J. et al. Accurate structure prediction of biomolecular interactions with AlphaFold 3. Nature 630, 493–500, doi:10.1038/s41586-024-07487-w (2024).

74 Elhan, H. et al. ATG2A-mediated DAG transfer recruits DGAT2 for lipid droplet growth. Nat Struct Mol Biol 32, 2601–2613, doi:10.1038/s41594-025-01689-0 (2025).

75 Monteiro-Cardoso, V. F. et al. ORP5 controls the partitioning of phosphatidic acid between triacylglycerol and cardiolipin synthesis at mitochondria-ER-lipid droplet contact sites. Preprint at bioRxiv 10.1101/2025.11.06.685814 (2025).

76 Ding, Y. et al. Isolating lipid droplets from multiple species. Nature Protocols 8, 43–51, doi:10.1038/nprot.2012.142 (2012).

77 Wieckowski, M. R., Giorgi, C., Lebiedzinska, M., Duszynski, J. & Pinton, P. Isolation of mitochondria-associated membranes and mitochondria from animal tissues and cells. Nature Protocols 4, 1582–1590, doi:10.1038/nprot.2009.151 (2009).

78 Lam, S. M. et al. Non-invasive lipid panel of MASLD fibrosis transition underscores the role of lipoprotein sulfatides in hepatic immunomodulation. Cell Metab 37, 69–86 e67, doi:10.1016/j.cmet.2024.09.009 (2025).

